# Genetic Analysis of *Arabidopsis* Autophagy-Related 8 Family Proteins Reveals Their Redundant and Regulatory Roles in Plant Autophagy

**DOI:** 10.1101/2025.01.01.631040

**Authors:** Kuntian Dong, Guiling Deng, Yilin Liu, Huan Wei, Kuo-En Chen, Xiner Huang, Wanying Huang, Ping Zheng, Takashi Ueda, Richard D. Vierstra, Xiao Huang, Faqiang Li

## Abstract

Autophagy, a critical process for the vacuolar degradation of proteins and organelles, is governed by multiple conserved autophagy-related (ATG) proteins. The central component of the ATG machinery is the ubiquitin-like protein ATG8, which is essential for multiple steps of the autophagy process, including phagophore expansion, autophagosome closure, trafficking and fusion with the lysosome/vacuole, and selective cargo recruitment. Currently, our understanding of the roles of ATG8 in plant autophagy and the functional specialization of ATG8 family members is limited due to genetic redundancy. To assess the roles of *ATG8* genes in plant autophagy, here we used CRISPR/Cas9 technology to systematically knockout the *Arabidopsis ATG8* genes. By analyzing the *atg8* mutants, we found that in contrast to mammalian ATG8s, in which the LC3s and GABARAP subfamilies play distinct roles in the autophagic process, *Arabidopsis* ATG8s perform an overlapping function in controlling autophagic flux. Combinatorial mutations of Clade I and Clade II ATG8s resulted in severely impaired autophagy under nutrient-starved conditions. Furthermore, we found that RABG3 proteins, members of the RAB7/RABG GTPase family, interact with ATG8s through AIM-LDS interfaces, and that such interaction is essential for the association of RABG3 proteins with the autophagosomal membrane and probably for the fusion of autophagosome with the vacuole, but is not required for endosomal trafficking. With the collection of multiple high-order *atg8* mutants generated in this study, we now provide a venue to study the roles of *ATG8* genes in canonical autophagy and non-canonical autophagy in *Arabidopsis*.

## Introduction

Autophagy is an intracellular catabolic pathway which plays critical roles in maintaining cellular homeostasis and helping cells to overcome nutrient scarcity. During autophagy, cytoplasmic materials are engulfed into double-membrane vesicles called autophagosomes, and targeted to the lysosomes (metazoans) or the vacuoles (yeast and plants) for degradation (Marshall & Vierstra, 2018; Nakatogawa, 2020; Zhao *et al*., 2021). Initially, autophagy was discovered as a non-selective bulk degradation process for survival under stress conditions. However, studies in the last two decades have revealed that autophagy is also a highly selective process by which distinct cellular components are recruited via dedicated receptors or adaptors (Johansen & Lamark, 2020).

When autophagy is induced, autophagic vesicles are generated at membrane cisternae called phagophores or isolation membranes, which are subsequently elongated and sealed by the orchestrated action of several conserved autophagy-related (ATG) proteins (Nakatogawa, 2020). The central component of the ATG machinery is the ubiquitin-like protein ATG8, which plays a vital role in all steps of the autophagic process, including phagophore expansion, autophagosome closure and trafficking, and fusion with lysosome/vacuole, as well as selective cargo recruitment (Kriegenburg *et al*., 2018). Like ubiquitin, ATG8 is firstly conjugated to lipid phosphatidylethanolamine (PE) at its carboxy-terminal glycine residue through a ubiquitin-like conjugation system, and then recruited to the inner and outer membranes of autophagic vesicles. The lipidated ATG8 proteins associated to the rim and outer surface of the phagophore facilitate phagophore growth by recruiting other autophagy core proteins (Xie *et al*., 2008; Nakatogawa, 2020). The outer membrane-anchored ATG8s are also involved in trafficking and vacuolar fusion of autophagosomes. For example, ATG8s bind to vesicle transport-related adaptor proteins, such as FYCO1 (FYVE and coiled-coil domain-containing 1) and JIP1 (JNK-interacting protein 1) to promote autophagosome trafficking (Pankiv *et al*., 2010; Fu *et al*., 2014), and bind to the core fusion machinery, including the MON1-CCZ1 complex, a guanine nucleotide exchange factor for Rab7, the SNARE protein STX17, and the tethering factor PLECKSTRIN HOMOLOGY DOMAIN-CONTAINING PROTEIN FAMILY MEMBER 1 (PLEKHM1) to coordinate the autophagosome-lysosome/vacuole fusion (Itakura *et al*., 2012; McEwan *et al*., 2015; Gao *et al*., 2018). In addition, ATG8s at the inner membrane act as a hub to recruit autophagic cargoes or receptors during selective autophagy. ATG8-interacting cargoes/receptors often contain one or multiple ATG8-interacting motifs/LC3-interacting regions (AIMs/LIRs) which recognize the LIR-docking site (LDS) in ATG8 proteins, or ubiquitin-interacting motifs (UIMs) which bind to the UIM-docking site (UDS) in ATG8s (Marshall *et al*., 2019; Johansen & Lamark, 2020).

In addition to the essential roles that ATG8s play in the autophagic process, which are known as its canonical functions, a growing number of studies in mammalian cells are revealing that ATG8s are involved in multiple alternative pathways that do not involve autophagosome formation (Nieto-Torres *et al*., 2021). During these so-called non-canonical autophagic processes, ATG8s are incorporated into divergent single-membrane vesicles with degradative or secretory functions, which are independent of the autophagy initiation machinery involved in canonical autophagy, but require the same ubiquitin-like conjugation system for lipidation. In plants, recent studies have begun to uncover the non-canonical functions of ATG8s. These studies found that ATG8s are translocated to swollen Golgi cisternae to aid in their reassembly after heat stress (Zhou *et al*., 2023), and to the tonoplast to maintain the vacuolar integrity after cell wall damage (Julian *et al*., 2024; Zheng *et al*., 2024; preprint). In addition, ATG8s were found to directly interact with the late endosome-resident transporter ABS3 to facilitate its vacuolar degradation in a non-canonical, lipidation-independent manner (Jia *et al*., 2019).

Although highly conserved across eukaryotes, ATG8 has diversified from a single protein in fungi and algae to multiple isoforms in mammals and higher plants (Kellner *et al*., 2017). At least eight ATG8 isoforms have been identified in mammals, which can be divided into two subfamilies (LC3 and GABARAP) based on their amino acid sequence homology. Genetic characterization and interactome studies revealed certain functional specialization for these two ATG8 subfamilies, as the LC3 subfamily is involved in the elongation of phagophore membrane and cargo recruitment, while GABARAP subfamily is essential for a later stage in autophagosome maturation and autophagosome-lysosome fusion (Weidberg *et al*., 2010; Johansen & Lamark, 2020). Similarly, the ATG8s in higher plants are often encoded by small gene families. For example, *Arabidopsis* and rice (*Oryza sativa*) have nine and five ATG8 isoforms, respectively. The nine isoforms (AtATG8a to AtATG8i) in *Arabidopsis* can be grouped into two major clades by phylogenetic analysis. Clade I includes AtATG8a-AtATG8g, and is closely related to yeast and other fungal homologs, whereas AtATG8h and AtATG8i, two isoforms with exposed Glycine residues at their carboxy termini that can be readily lipidated, form clade II together with animal ATG8 homologs (Kellner *et al*., 2017). However, whether the two *Arabidopsis* ATG8 clades are functionally specialized in plant autophagy biogenesis remains to be elucidated.

Currently, our understanding of the roles of ATG8s in plant autophagy and the functional specialization of ATG8 family members is limited due to genetic redundancy. Numerous genetic analyses in *Arabidopsis* and other plants have suggested that ATG8s are important for plant growth and development, and for survival under various stress conditions, mostly based on the results of homozygous null mutants defective in ATG8 lipidation or from plants overexpressing ATG8s (Doelling *et al*., 2002; Thompson *et al*., 2005; Phillips *et al*., 2008; Chung *et al*., 2010; Xia *et al*., 2012; Wang *et al*., 2016; Chen *et al*., 2019; Fan *et al*., 2020; Zhen *et al*., 2021; Kanne *et al*., 2022). Recently, high-order mutants have been used to study ATG8 functions. For example, Lan *et al*. (2024) reported that simultaneous knockout of *Arabidopsis* ATG8h and ATG8i lead to enhanced resistance against the biotrophic fungal pathogen *Golovinomyces cichoracearum*, likely through a specific interaction with CLATHRIN LIGHT CHAIN (CLC) subunit 2 and 3. Simultaneous knockout of multiple rice ATG8s resulted in delayed flowering, likely due to a defect in autophagy-mediated degradation of the central regulator of flowering Hd1 (Hu *et al*., 2022). In addition, by characterizing ATG8 interactors, including SH3 DOMAIN-CONTAINING PROTEIN2 (SH3P2) (Zhuang *et al*., 2013; Sun *et al*., 2022), FYVE DOMAIN PROTEIN REQUIRED FOR ENDOSOMAL SORTING1 (FREE1)/FYVE1 (Zeng *et al*., 2023) and FYVE2 (also known as CFS1) (Kim *et al*., 2022; Zhao *et al*., 2022), and Sar1d (Zeng *et al*., 2021), the roles of ATG8s in plant phagophore expansion, autophagosome closure and maturation are beginning to be elucidated, strengthening the notion that ATG8 is a key player in the plant autophagic process.

The Rab GTPase RAB7 and its yeast counterpart Ypt7 are essential for endocytic membrane trafficking from late endosome to lysosome/vacuole, and for autophagosome-lysosome fusion (Guerra & Bucci, 2016). RAB7/Ypt7 targeting to autophagosomes and its activation are triggered by the ATG8-binding guanine nucleotide exchange factor MON1-CCZ1 complex (Hegedűs *et al*., 2016; Gao *et al*., 2018). Subsequently, RAB7 interacts with multiple downstream effectors, including tethering factors such as HOPS (Homotypic fusion and protein sorting) complex, PLEKHM1, and EPG5, to drive membrane fusion (Zhao *et al*., 2021). Similarly, RAB7 GTPases function as coordinators of plant endomembrane traffic and are essential for late endosome-tonoplast fusion (Rodriguez-Furlan *et al*., 2023). In *Arabidopsis*, RAB7 is recruited to prevacuolar compartments and activated by the MON1-CCZ1 complex to regulate vacuolar trafficking, vacuole biogenesis, and plant growth (Cui *et al*., 2014; Singh *et al*., 2014). However, the precise role of RAB7 in plant autophagy and how it is recruited to the autophagosomal membrane is still unclear.

To investigate the roles of ATG8 genes in plant autophagy, here we used CRISPR/Cas9 technology to systematically mutate the *Arabidopsis ATG8* genes. By analyzing the *atg8* mutants, we found that in contrast to mammalian ATG8s, in which the LC3s and GABARAP subfamilies act in the early and late stages of the autophagic process, respectively, the *Arabidopsis ATG8* genes have an overlapping function in controlling autophagic flux. Combinatorial mutations of clade I and clade II ATG8s showed severely impaired autophagy under nutrient-starved conditions. Furthermore, we found that RABG3 proteins, members of the RAB7/RABG GTPase family, interact with ATG8s through AIM-LDS interfaces, and that such interaction is essential for the association of RABG3 proteins with autophagosomes and likely for autophagosome-vacuole fusion, but not for endosomal trafficking pathways. With the collection of several high-order *atg8* mutants generated in this study, we now provide a venue to study the roles of *ATG8* genes in canonical autophagy and non-canonical autophagy in *Arabidopsis*.

## METHODS

### Plasmid Construction

For the constructs used for transient expression in *Arabidopsis* protoplasts, the coding sequences (CDS) of *SH3P2*, *NBR1*, *ABS3*, *RABG3f*, and several *ATG* genes (*ATG1a, ATG9*, *ATG14a*, *ATG5*, and *ATG8a*) were amplified using the oligonucleotides described in Supplemental Table 1 and cloned into pBI221 vectors modified to contain GFP, or mCherry tags driven by the 35S promoter (Li *et al*., 2022). The pBI221-mCherry-RABG3f^mAIM1,2^ construct was generated by PCR mutagenesis to introduce point mutations (F94A/L99A/F167A/I172A) in the AIM sequences of RABG3f, and cloned into pBI221 vector with mCherry tag. The Aleu-GFP construct has been reported previously (Cui *et al*., 2017).

To generate the constructs for investigating RABG3-ATG8 protein interactions in tobacco (*Nicotiana benthamiana*) using the LCI assay, the CDS of *RABG3f*, *RABG3b*, *RABG3d*, *RABG3e*, *ATG8a*, *ATG8e* and *ATG8h* were amplified and cloned into pCAMBIA1300-nLUC and pCAMBIA1300-cLUC, respectively (Chen *et al*., 2008). The constitutively active (CA, Q67L), dominant negative (DN, T22N) and various AIM mutated variants of RABG3f, RABG3f(CA), RABG3f(DN), RABG3f^mAIM1^, RABG3f^mAIM2^, RABG3f^mAIM1,2^, were generated from RABG3f-nLUC by PCR mutagenesis.

To generate transgenic plants coexpressing GFP-ATG8a and mCherry-RABG3f or mCherry-RABG3f^mAIM1,2^, their full-length coding sequences were inserted into a modified pCAMBIA1300 vector carrying two different expression cassettes for in-frame fusion with a GFP or mCherry tag at the N-terminus driven by the constitutive *AtUBQ10* promoter. The resulting constructs (pUBQ10:mCherry-RABG3f-NOS-pUBQ10:GFP-ATG8a-NOS and pUBQ10:mCherry-RABG3f^mAIM1,2^-NOS-pUBQ10:GFP-ATG8a-NOS) were transformed into *Agrobacterium tumefactions* GV3101 and then introduced into *rabg3a,b,c,d,e,f* (*rabg3f-6m*, Ebine *et al*., 2014), mutant plants using the floral dip method (Clough & Bent, 1998). Homozygous *rabg3-6m* plants carrying the transgene were identified in the T3 generation based on hygromycin resistance and confocal fluorescence microscopy.

### Plant Materials and Growth Conditions

The *Arabidopsis thaliana* accession Col-0 was used as the wild type in this study. The T-DNA mutants *atg5-1* (Thompson *et al*., 2005), *atg7-2* (Chung *et al*., 2010), *gfs9-3* (Ichino *et al*., 2014) and *rabg3-6m* (Ebine *et al*., 2014), as well as transgenic plants expressing p35S:GFP-ATG8a (Thompson *et al*., 2005), ProUBQ10:GFP-ATG8a (Shin *et al*., 2014), pUBQ10:mCherry-RabG3f (Geldner *et al*., 2009), all in the Col-0 background, have been described previously. *gfs9-4/tt9* mutant (Ichino *et al*., 2014) is in the Landsberg *erecta* (L*er*) background. Fluorescent protein expression cassettes p35S:GFP-ATG8a and pUBQ10:mCherry-RabG3f were introgressed into the *gfs9-4/tt9* mutant by crossing. Segregants containing the Col-0 allele of *GFS9* were used as control lines. To determine the effect of the RABG3 mutation on autophagic activity, a construct carrying ProUBQ10:GFP-ATG8a (hygromycin resistance, Shin *et al*., 2014) was transformed into the *rabg3-6m* lines by the floral dip method. Homozygous *rabg3-6m* plants carrying the transgene were identified in the T3 generation based on hygromycin resistance and confocal fluorescence microscopy.

Unless otherwise indicated, all *Arabidopsis* seeds were surface sterilized using vapor-phase method and stratified in water at 4°C for two days in the dark before sowing on Murashige and Skoog (MS) solid medium supplemented with 1% (w/v) sucrose (Suc). Plants were grown at 22°C under long-day (LD, 16-h light/8-h dark) conditions for ten days before being transferred to soil for further growth. For senescence assays, plants were grown in soil under LD conditions for seven weeks or under a short-day (SD, 8-h light/16-h dark) photoperiod at 22°C for ten weeks.

For N starvation treatment, seven-day-old seedlings grown on MS solid medium with 1% (w/v) Suc were transferred to MS liquid medium with or without nitrogen. Seedlings were grown under continuous white light irradiation for the indicated times before imaging and measuring chlorophyll content as previously described (Suttangkakul *et al*., 2011). Alternatively, seeds were germinated on MS solid medium containing 1% (w/v) Suc and 1.4% (w/v) agar with or without a nitrogen supply and grown vertically. After seven days, seedlings were imaged and root lengths were determined as described previously (Xiao *et al*., 2020).

For carbon starvation, two-week-old seedlings grown on MS solid medium without Suc were wrapped in aluminum foil and kept in the dark for the indicated times before recovery under LD conditions. After two weeks of recovery, seedlings were imaged and survival rates were determined as previously described (Suttangkakul *et al*., 2011). Alternatively, seven-day-old seedlings grown on MS solid medium containing 1% (w/v) Suc of uniform size were selected and transferred to MS solid medium without Suc, and grown vertically under continuous darkness for the indicated time before imaging and measuring chlorophyll content.

### Generation of the *atg8* Nonuple Mutants

The constructs for CRISPR were generated by using an egg cell-specific promoter-controlled CRISPR/Cas9 genome editing system (Xing *et al*., 2014; Wang *et al*., 2015). We constructed three CRISPR/Cas9 vectors to knock out *ATG8abcd*, *ATG8efg* and *ATG8hi*, respectively. Briefly, the guide RNAs targeting different *ATG8* genes were designed by using the CRISPR-GE website (http://skl.scau.edu.cn/) and cloned into the pHEE401E vector. The cloning primers, the arrangement of the sgRNAs in three CRISPR/Cas9 constructs and the sequences of the constructed plasmids are provided in the supplementary materials (Supplemental Table 1 and Supplemental Doc). The generated constructs were then introduced into the wild-type Col-0 plants by the Agrobacterium-mediated floral-dip method. The resulting T1 transformants were selected for hygromycin resistance and positive plants were transferred to soil after two weeks. Mutations were identified by PCR amplification of the genomic regions encompassing the DNA target sites and sequencing (See Supplemental Table 1 for PCR primers). In the next generation, mutants were further examined by PCR and sequencing to obtain homozygous mutants without hygromycin resistance and the corresponding CRISPR/Cas9 vector. After obtaining hygromycin-sensitive Cas9-free homozygous mutants, CRISPR/Cas9 constructs targeting *ATG8abcd* and *ATG8hi* were introduced into the *atg8efg* triple mutant (abbreviated as *atg8-3m*) to generate *atg8efghi* quintuple mutant (*atg8-5m*) and *atg8abcdefg* septuple mutant (*atg8-7m*), respectively. After obtaining a hygromycin-sensitive Cas9-free homozygous *atg8-7m* mutant, the CRISPR/Cas9 construct targeting *ATG8hi* was further transferred into the mutant to generate *atg8abcdefghi* nonuple mutants (*atg8-9m*).

### Transient Expression in *Arabidopsis* Protoplasts

The *Arabidopsis* leaf protoplast preparation and transient expression were performed according to a standard procedure (Yoo *et al*., 2007). Briefly, confocal images were collected at 12 to 16 h or at the indicated time using a Leica Stellaris 5 confocal microscope (Leica, Wetzlar, Germany). To detect autophagic bodies in protoplasts, 1 μM ConA (AdipoGen life sciences, BVT-0237-M001) or an equivalent volume of DMSO was added to the protoplasts for 12 h prior to observation.

### Fluorescence Confocal Microscope Imaging and Analysis

Fluorescence cell images were collected using a Leica Stellaris 5 confocal microscope. Confocal imaging of stable transgenic lines expressing GFP-ATG8a in wild type, and *rabg3-6m* and *gfs9-4* mutant backgrounds was performed as described previously (Suttangkakul *et al*., 2011; Huang *et al*., 2019). Briefly, six-day-old seedlings grown on nitrogen-containing MS solid medium were transferred to either fresh nitrogen-rich medium or nitrogen-deficient medium supplemented with 1 μM concanamycin A (ConA) for the indicated time, and confocal images of the root cell were captured. To acquire the GFP signal, a 488-nm laser was used for excitation, and fluorescence was detected in the range of 490-540 nm. To image the coexpression of GFP and mCherry constructs in leaf protoplasts and the root cells of stable transgenic lines, excitation wavelengths of 488 nm for GFP and 543 nm for mCherry were used alternately in the multitrack mode of the microscope with line switching. Quantitative analysis of confocal microscopy images was performed by using ImageJ (https://imagej.nih.gov/) as described by Kim *et al*. (2022).

### Luciferase Complementation Imaging (LCI) Assay

To investigate the protein interaction between RABG3 and ATG8, the LCI assays were performed as described previously (Chen *et al*., 2008). Briefly, different combinations of RABG3 and ATG8 constructs or empty vectors were introduced into *Agrobacterium tumefaciens* strain GV3101 and then co-infiltrated into tobacco leaves. After infiltration, the plants were immediately cultured in the dark for 16 h and then exposed to LD conditions for additional 48 h at 23°C. The agro-infiltrated leaves were further infiltrated with 500 µM D-luciferin, and luminescence signals were captured using a low-light cooled charge-coupled device camera (Night owl LB985, Berthold Technologies, Germany). After observation, the infiltrated leaves were collected for Western blot to examine the expression levels of nLUC and cLUC fusion proteins.

### GST Pull-Down Assay

GST-ATG8e was produced in *Escherichia coli* BL21 (DE3) after induction with 1 mM IPTG at 16°C for 16 h. Total *E. coli* proteins were then extracted and incubated with Glutathione Sepharose beads (C650031, Sangon Biotech Co., Ltd., China) for GST-ATG8e purification. For the pull-down analysis, the GFP-nLUC, RABG3f-nLUC, and RABG3f-nLUC^mAIM1,2^ were individually expressed in tobacco leaves by agroinfiltration approach. After three days, approximately 500 mg infiltrated leaves were harvested and homogenized with 2 mL lysis buffer (50 mM Tris-HCl, pH 7.4, 150 mM NaCl, 1 mM MgCl2, 20% glycerol, 0.2% NP-40, and 1X protease inhibitor cocktail from Roche). Cell lysates were then clarified twice by centrifugation at 13,000 g for ten min at 4°C and were incubated with GST-ATG8e-bound beads for one h at 4°C. The beads were then washed five times with 1 mL of wash buffer (140 mM NaCl, 2.7 mM KCl, 10 mM Na_2_HPO4, 1.8 mM KH_2_PO4, pH 7.4) for one min each time. Protein bound to the beads were released in SDS-PAGE loading buffer by boiling for five min and subjected to immunoblot analysis.

### Protein Extraction and Immunoblot Assay

To detect different ATG proteins and GFP/mCherry-tagged proteins by immunoblot assays, 100 mg of seedlings were homogenized in 200 mL of 2X SDS-PAGE sample buffer (100 mM Tris-HCl, pH 6.8, 4% [w/v] SDS, 0.2% [w/v] bromophenol blue, and 20% [v/v] glycerol), followed by heating at 95°C for five min. The plant lysates were then clarified twice by brief centrifugation at 14,000g for ten min at room temperature and the resulting supernatants were subjected to SDS-PAGE and immunoblot analysis. Specific anti-ATG1a (Suttangkakul *et al*., 2011), anti-ATG5 and anti-ATG8a (Thompson *et al*., 2005), anti-PBA1 (Smalle *et al*., 2002), anti-GFP (ab290; Abcam, Cambridge, UK), anti-mCherry (B1153; Biodragon, Beijing, China), anti-GST (B1007; Biodragon) and anti-luciferase (L0159, Sigma-Aldrich) antibodies were used for protein blot analysis. Protein levels were quantified using ImageJ software as previously described (Kim *et al*., 2022).

### Protein Sample Preparation for MS Analysis

To prepare plant samples for MS analysis, sterilized wild-type and *atg* mutant seeds were grown under continuous light in 250 mL flasks containing 50 mL of MS liquid medium supplemented with 1% Suc for seven days. For nitrogen starvation, the plants were transferred to MS liquid medium with or without nitrogen for three days. For carbon starvation, the plants were transferred to MS liquid medium with or without Suc for two days, and the flasks containing carbon-starved plants were covered with aluminum foil. Seedlings were then harvested and homogenized in liquid nitrogen and extraction buffer (50 mM HEPES, 5 mM EDTA, 2 mM DTT, and 1X protease inhibitor cocktail). Total extracted proteins were precipitated using a 4:1:3 (v/v) methanol/chloroform/water mixture, collected by centrifugation, washed once with the methanol, and then lyophilized to dryness. The precipitates were then resuspended in 100 µL of 8 M urea and reduced with 10 mM dithiothreitol for 1 hour at room temperature. For trypsin digestion, 100 ug of protein was alkylated with 20 mM iodoacetamide for 1 hour, and the reaction was quenched with 20 mM dithiothreitol. The sample was diluted with 900 µL of 25 mM ammonium bicarbonate to reduce the urea concentration to below 1 M, and digested overnight at 37°C with sequencing grade modified porcine trypsin (Promega) at a trypsin:protein ratio of 1:50. The resulting peptides were lyophilized to a volume of less than 50 µL, acidified with 10% trifluoroacetic acid to a pH below 3.0, and desalted and concentrated using Pierce C18 tips (Thermo Fisher Scientific) according to the manufacturer’s instructions. Peptides were eluted in 50 µL of 75% acetonitrile and 0.1% acetic acid, lyophilized again, and resuspended in 15 µL of 5% acetonitrile and 0.1% formic acid for LC-MS/MS analysis.

### Mass Spectrometric Analysis and Data Process

Full MS scans were performed in the mass range of 380-1500 m/z at a resolution of 70,000, with an automatic gain control target of 3 × 10^6^ and a maximum injection time of 200 ms. Data-dependent acquisition (DDA) was applied to fragment the top fifteen most intense peaks using high-energy collision-induced dissociation (HCD) at a normalized collision energy of 28. An intensity threshold of 4 × 10^4^ counts and an isolation window of 3.0 m/z were applied. Precursor ions with unassigned charges or charges between +1 and +7 were excluded. MS/MS scans were collected at a resolution of 17,500, with an AGC target of 2 × 10^5^ and a maximum fill time of 300 ms. Dynamic exclusion was performed with a repeat count of 2 and an exclusion duration of 30 s, while the minimum MS ion count to trigger tandem MS was set to 4 × 10^3^ counts. Raw sequencing data are deposited in the PRoteomics IDEntifications Database.

The resulting MS/MS spectra were analyzed using Proteome Discoverer (version 2.5, Thermo Fisher Scientific) against the *A*. *thaliana* Col-0 proteome database (Araport11_pep_20220914), downloaded from TAIR (http://www.tair.com/). Peptide assignments were performed using SEQUEST HT with the following parameters: trypsin digestion allowing a maximum of two missed cleavages, a minimum peptide length of six residues, a precursor mass tolerance of 10 ppm, and a fragment mass tolerance of 0.02 Da. Carbamidomethylation of Cys and oxidation of Met were specified as static and dynamic modifications, respectively. False discovery rates of 0.01 (high confidence) and 0.05 (medium confidence) were used to validate peptide spectral matches. Label-free quantification based on MS1 precursor ion intensity was performed in Proteome Discoverer with a minimum Quan value threshold set to 0.0001 for unique peptides; the “3 Top N” peptides were used for area calculation. All genotypes and treatments were analyzed in four biological replicates, each with two technical replicates.

Protein abundances were normalized using the values of the 150 least variable proteins (determined by the standard deviation/average ratio) across samples for each run. Protein abundances were then further adjusted based on protein content per weight of plant tissue for each biological replicate. The normalization was validated by comparing the abundances of the detected tubulin proteins (Fig. 3A and Supplemental Figure 7). Standardized data were used to filter proteins that must be present in all four biological replicates under at least one genotype. To generate volcano plots, data were first transformed to Log_2_ values, with missing value imputation and statistical calculations conducted in Perseus software, followed by visualization using GraphPad Prism (version 10). Statistic differences between *atg* mutants and WT were determined based on four biological replicates using Student’s *t*-test (Log_2_ FC ≥ 1 or ≤ –1, *P* ≤ 0.05). GO analyses were performed using the *Arabidopsis* profile database in g-:Profiler V3.10.178 as part of the ELIXIR Infrastructure package (http://biit.cs.ut.ee). The GO-annotation categories shown here were selected based on their uniqueness, *P*-values of significance, and degrees of completeness.

## RESULTS

### Multigene Knockouts of *Arabidopsis ATG8* Genes

The *Arabidopsis* ATG8 protein family consists of nine members (ATG8a to ATG8i), which can be divided into two major clades based on their sequence homology and genomic structure (Kellner *et al*., 2017). Clade II contains ATG8h and ATG8i isoforms, while clade I can be further split into two subclades (I-1, ATG8a-d and I-2, ATG8e-g). To investigate the significance of *Arabidopsis* ATG8s in the autophagic process, we used an egg cell-targeting clustered regulatory interspaced short palindromic repeats (CRISPR)/Cas9 technique to systematically edit the *ATG8* genes (Wang *et al*., 2015). Three CRISPR/Cas9 vectors were constructed and targeted *ATG8* genes within subclade I-1 (*ATG8a-d*), subclade I-2 (*ATG8e-g*), and clade II (*ATG8h-i*), respectively (Figure. 1A and B, and Supplemental Doc). Through *Agrobacterium*-mediated transformation of the *Arabidopsis* wild-type Col-0, first generation seedlings with specific mutations in *ATG8s* were obtained. Next, we used the CRISPR/Cas9 vectors to further knock out *ATG8a-d* and *ATG8h-i*, respectively, in the Cas9-free *atg8efg* triple mutant (abbreviated as *atg8-3m*) to obtain *atg8efghi* quintuple mutant (abbreviated as *atg8-5m*) and *atg8abcdefg* septuple mutant (abbreviated as *atg8-7m*), respectively. Finally, we generated an *atg8abcdefghi* nonuple mutant (abbreviated as *atg8-9m*) in the cas9-free *atg8-7m* background using CRISPR/Cas9 to knock out *ATG8h* and *ATG8i*. Mutations in double (*atg8hi*, also referred to as *atg8-2m*), triple, quadruple (*atg8abcd*, also referred to as *atg8-4m*), quintuple, septuple, and nonuple were confirmed by PCR and DNA sequencing as shown in Supplemental Figure S1.

**Figure 1.**
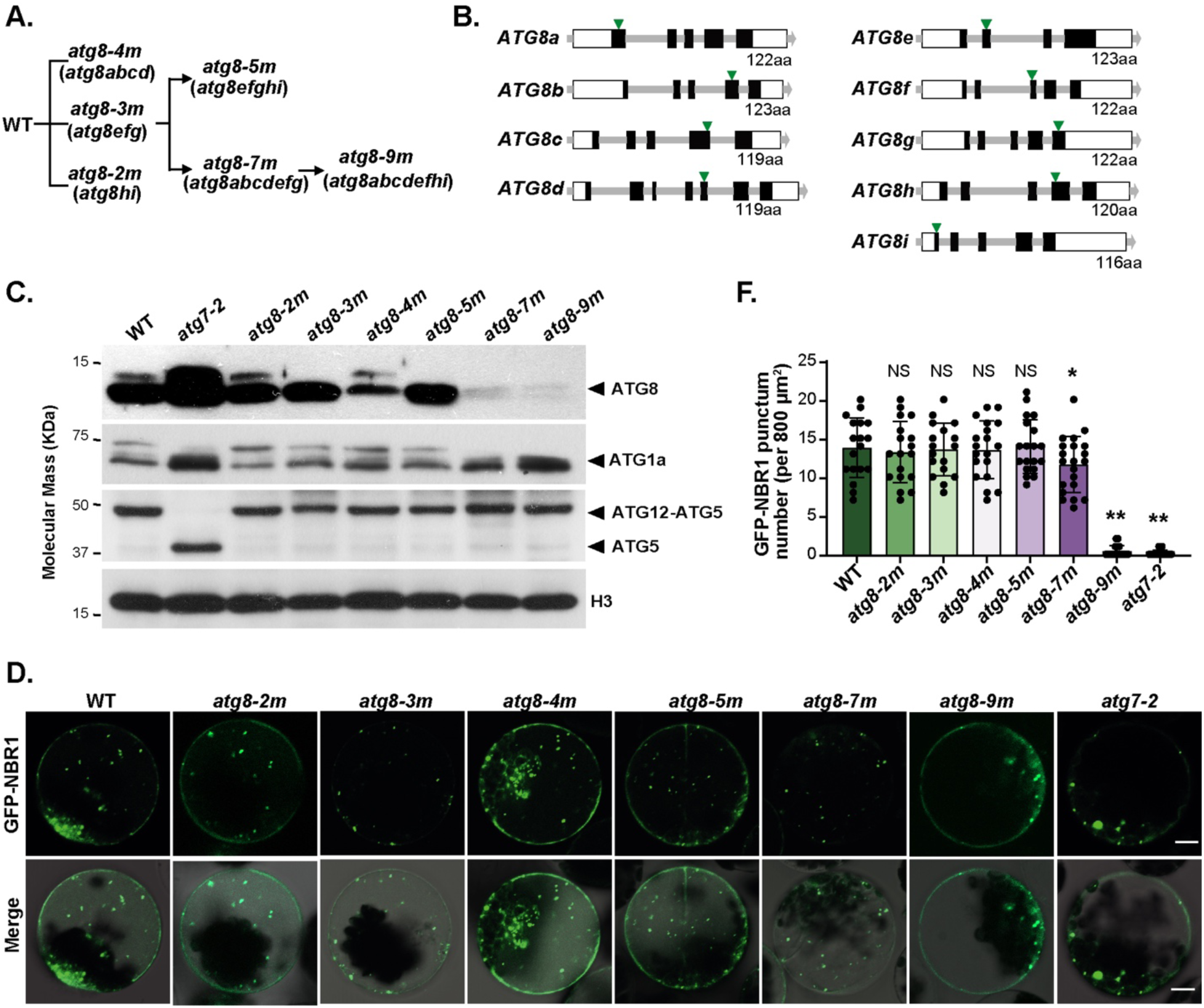
Generation of *atg8* Nonuple Mutants. **(A)** Generation of *atg8* nonuple mutants. Three CRISPR/Cas9 constructs targeting *ATG8abcd*, *ATG8efg*, or *ATG8hi* were constructed and transformed into wild-type Col-0 to generate *atg8abcd* quadruple mutant (*atg8-4m*), *atg8efg* triple mutant (*atg8-3m*), and *atg8hi* double mutant (*atg8-2m*), respectively. Subsequently CRISPR/Cas9 constructs targeting *ATG8abcd* and *ATG8hi* were transferred into the *atg8-3m* mutant to generate the *atg8efghi* quintuple mutant (*atg8-5m*) and the *atg8abcdefg* septuple mutant (*atg8-7m*), respectively. To generate *atg8* nonuple mutants, a CRISPR/Cas9 construct targeting *ATG8hi* was transferred into the *atg8-7m* mutant. **(B)** Diagram illustrating the *ATG8* genes and the selected target sites. Exons are shown as black boxes, 5′-UTR and 3′-UTR as white boxes, and sgRNA target sites as green triangles. **(C)** Immunoblot detection of multiple ATG proteins from wild-type (WT), *atg7-2*, and various *atg8* mutants using antibodies against ATG8, ATG1a, and ATG5, respectively. Antibody against H3 was used as a loading control. **(D)** Effects of different *atg8* mutations on the vacuolar transport of GFP-NBR1. Leaf protoplasts of WT, *atg7-2*, or various *atg8* mutants were transformed with the GFP-NBR1 construct and treated with 1 μM ConA for 12 to 14 h before confocal imaging analysis. Bars = 10 μm. **(E)** Quantification of the number of vacuolar GFP-NBR1 puncta per 800 μm^2^, using images (*n* = 18-22) similar to those shown in (D). Asterisked columns represent *atg8* mutants that are significantly different from WT, according to Student’s *t*-test. *0.01<*P* <0.05; \*\**P* < 0.01. NS, not significant.

To investigate how different *atg8* mutations affect the accumulation of ATG8 proteins, we performed immunoblot analysis using antibodies raised against the recombinant *Arabidopsis* ATG8a isoform, which also readily identified the most divergent ATG8 isoform, ATG8i (Doelling *et al*., 2002). As shown in Figure 1C, ATG8 levels remained high in the seedlings of *atg8-2m*, *atg8-3m*, *atg8-4m*, and *atg8-5m* mutants, whereas they were significantly reduced in homozygous *atg8-7m* lines. In *atg8-9m* seedlings, only trace amounts of ATG8 proteins were detected, which most likely represent the truncated forms of ATG8 isoforms, such as ATG8g, generated by CRISPR/Cas9 editing (Figure 1B and 1C). Previous studies with ATG1 suggested that it is not only a regulator but also a cargo of autophagic degradation (Suttangkakul *et al*., 2011). In *atg8-7m* and *atg8-9m* seedlings, ATG1 levels were constitutively upregulated as compared with wild type, similar to that observed for the *atg7-2* mutant, suggesting that ATG1 degradation was impaired in both *atg8-7m* and *atg8-9m* mutants (Figure 1C). In contrast, the ATG12–ATG5 conjugate required for ATG8 lipidation was readily detected in all *atg8* mutants using anti-ATG5 antibodies, as seen in the wild type. Notably, the amount of free ATG5 increased subtly in *atg8-9m* seedlings (Figure 1C, see also Figure 6F), suggesting that a complete removal of *ATG8* genes somehow affects the assembly of ATG12–ATG5.

In addition to immunoblot analysis, we also used protoplast transient assay to investigate whether the ATG8 mutations affect the subcellular localizations of other ATG proteins, the selective autophagy receptor NBR1 and the MATE transporter ABS3. In contrast to the wild-type protoplast, no GFP-ATG1a and GFP-ATG14 puncta were found in *atg7-2* or *atg8-9m* cells treated with the specific v-ATPase inhibitor ConA (Supplemental Figure 2A and 2B), consistent with the notion that both ATG1 and ATG14 are delivered to the vacuole through their association with autophagic vesicles (Suttangkakul *et al*., 2011; Liu *et al*., 2020). On the other hand, the subcellular distributions of other ATG components, including ATG9, SH3P2, and ATG5, were unperturbed in *atg7-2* or *atg8-9m* protoplasts compared with the wild type (Supplemental Figure 2C). For AtNBR1-GFP, its fluorescence appeared as punctate structures in the cytosols of all mutant protoplasts, including *atg7-2* and *atg8-9m* cells before ConA treatment (Supplemental Figure 3). However, after ConA treatment, NBR1-GFP puncta were readily detected in the vacuoles of wild-type, *atg8-2m*, *atg8-3m*, *atg8-4m*, *atg8-5m* and *atg8-7m* cells, but not in *atg7-2* and *atg8-9m* mutants (Figure 1D). For ABS3-GFP, we detected endosomal ABS3-GFP puncta in all DMSO-treated protoplasts and found punctuate ABS3-GFP signals in the vacuoles of ConA-treated wild-type and *atg7-2* protoplasts (Supplemental Figure 4), consistent with the previous study that the vacuolar trafficking of ABS3-GFP is independent of the ATG8 conjugation machinery (Jia *et al*., 2019). However, the deposition of ABS3-GFP in the vacuole was completely abolished in the *atg8-9m* background, as ABS3-GFP maintained a predominantly endosomal distribution in the cell periphery (Supplemental Figure 4). Taken together, these results demonstrate that *atg8-9m* represents a functionally null mutant in which both non-selective autophagy and selective autophagy, as well as autophagy-independent ABS3 proteolysis, are severely impaired.

**Figure 2.**
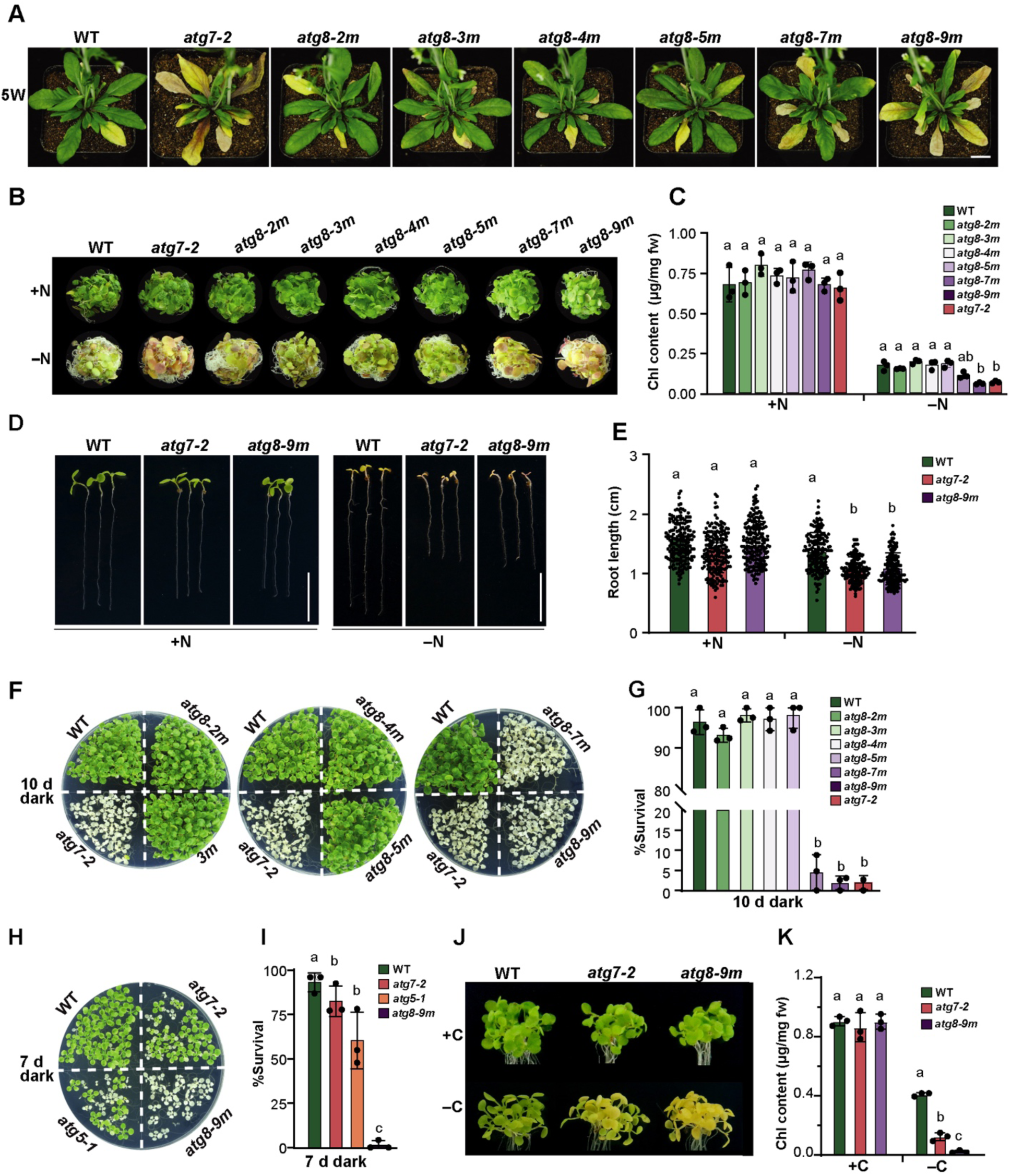
Phenotypic Analyses of *Arabidopsis atg8* Mutants. The homozygous *atg8* mutants described in Figure 1, the autophagy mutants *atg5-1* and *atg7-2*, and the wild-type Col-0 (WT) were included for comparisons. **(A)** Accelerated senescence. Plants were grown on soil at 22°C under a long-day (LD) photoperiod (16-h-light/8-h-dark) for five weeks. Bar = 1 cm. **(B)** Enhanced sensitivity to nitrogen starvation. Seeds were germinated in 1x MS liquid medium and grown under constant white light conditions for one week, and then transferred to either fresh MS (+N) or nitrogen-deficient (–N) liquid medium for an additional one week. **(C)** Total chlorophyll content of the plants shown in (B). Data are presented as mean ± s.d. (*n* = 3 biological replicates, 80-120 seedlings per genotype in each independent experiment). **(D)** Short primary root phenotype of *atg8-9m* in response to nitrogen deficiency. Seeds were germinated and grown vertically on MS medium with (+N) or without nitrogen (–N) under LD conditions for one week. Bar = 1 cm. **(E)** Quantification of root length of the plants shown in (D). Data are presented as mean ± s.d. (*n* = 3 biological replicates, >50 seedlings per genotype in each independent experiment). **(F and H)** Enhanced sensitivity to fixed-carbon starvation. Seedlings were grown under an LD photoperiod on MS solid medium without sucrose (–C) for two weeks, transferred to darkness for 10 d (F) or 7 (H) d, and then allowed to recover under LD conditions for 12 d. **(G and I)** Quantification of the effects of fixed carbon starvation based on seedling survival after 10 (F) or 7 (H) d in darkness followed by 12 d in LD. Data are presented as mean ± s.d. (*n* = 3 biological replicates, 60-120 seedlings per genotype in each independent experiment). **(J)** Senescence phenotypes of the *atg8-9m* after dark treatment. Seedlings were germinated and grown on 1/2 MS solid medium with 1% sucrose under LD conditions for 1 week, then transferred to 1/2 MS medium without sucrose (–C) and grown vertically in the dark for another seven d. **(K)** Total chlorophyll content of the plants shown in (J). Data are presented as mean ± s.d. (*n* = 3 biological replicates, 80-120 seedlings per genotype in each independent experiment). Different letters in (C), (E), (G), (I), and (K) indicate significant differences (*P* < 0.05) as determined using two-way ANOVA followed by Tukey’s multiple comparison test.

**Figure 3.**
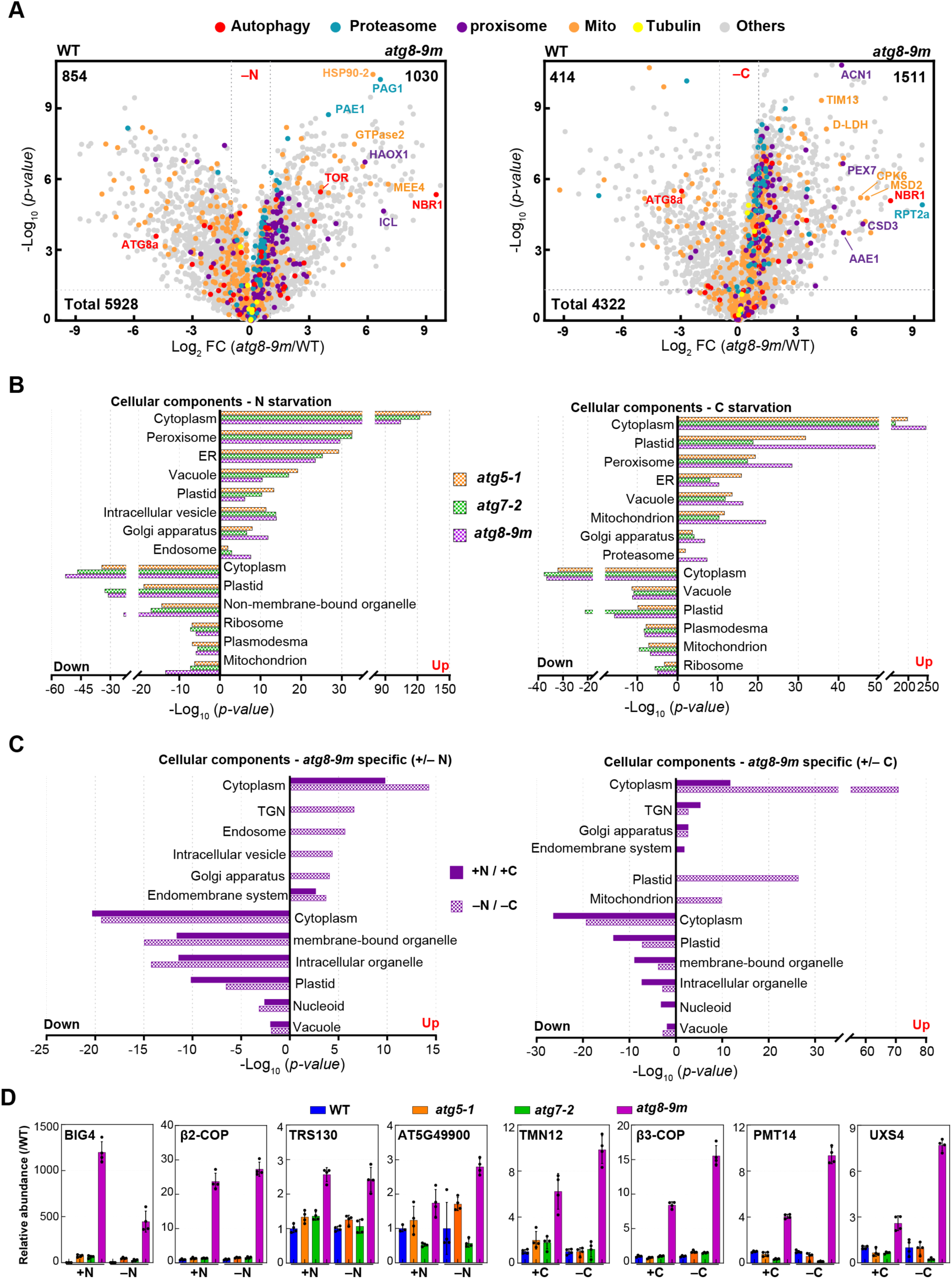
The *Arabidopsis* Proteome Is Strongly Affected by the *atg8-9m* Mutation. **(A)** Volcano plots showing the preferential accumulation of proteins categorized by Go to autophagy process, proteasome complex and specific cellular compartments in *atg8-9m* versus wild-type seedlings under N-or C-starvation. Protein abundances were measured using MS1 precursor ion intensities based on the average of four biological replicates each analyzed in duplicate. Proteins assigned to the autophagy, 26S proteasome, peroxisome, and mitochondrion are colored in red, cyan, purple, and orange, respectively. Detected tubulins and proteins within other GO categories are labeled in yellow and gray, respectively. **(B)** Log_10_ fold enrichment/depletion using a singular enrichment of specific GO terms for proteins that were consistently altered in abundance in the *atg* mutants compared with the wild type (+/– N or +/–C). **(C)** Specific GO terms for proteins that were significantly enriched or depleted only in the *atg8-9m* plants. **(D)** Levels of representative Golgi proteins specifically affected by the *atg8-9m* mutant. Each value was normalized to the average value of the wild type and presented as mean ± s.d. (*n* = 4 biological replicates, each analyzed in duplicate by MS).

### Plants Lacking All ATG8 Isoforms Senesce Early and Are Hypersensitive to Nutrient-Limiting Conditions

To determine how severely the *atg8* mutations compromise plant autophagy, we performed a series of phenotypic analyses as previously reported (Huang *et al*., 2019). Both ATG proteins, ATG5 and ATG7, have been shown to play essential roles in ATG8 lipidation, whereas plants lacking ATG5 or ATG7 are characterized by their early senescence phenotypes and hypersensitivity to nutrient deprivation (Thompson *et al*., 2005; Chung *et al*., 2010). Therefore, the null alleles *atg5-1* and *atg7-2* were included in this study as positive controls for phenotypic responses in the following analyses. As shown in Supplemental Figure 5, all *atg8* mutants developed and grew normally in nutrient-rich soil under greenhouse conditions. Furthermore, all *atg8* mutants produced viable pollen and were fully fertile under greenhouse conditions, as judged by vital staining with Alexander’s stain and *in vitro* pollen germination (Supplemental Figure 6). However, the *atg8-7m* and *atg8-9m* exhibited an early senescence phenotype compared with the wild type, phenocopying the *atg7-2* mutant (Figure 2A). Under long-day (LD) conditions, both *atg8-7m* and *atg8-9m* mutants began to senesce as early as five weeks after planting, approximately 1.5 weeks before the wild type and other *atg8* mutants.

Next, we challenged the *atg8* mutants with several nitrogen– and fixed carbon–limiting conditions to assess their sensitivity and survival. Similar to *atg7-2* seedlings, *atg8-9m* plants grew poorly and became more chlorotic when grown under nitrogen-starved conditions (Figure 2B and 2C). Notably, the *atg8-7m* mutant also showed a slight sensitivity to nitrogen starvation. In addition, the primary roots of *atg8-9m* seedlings showed shorter primary roots on nitrogen-limited medium (Figure 2D and 2E). For fixed-carbon starvation, both *atg8-7m* and *atg8-9m* plants were as hypersensitive as the *atg7-2* mutant when plants were subjected to ten days of darkness, and nearly all of these mutants died, compared with 4-7% mortality for wild-type plants and other *atg8* mutants (Figures 2F and 2G). Surprisingly, *atg8-9m* seedlings showed higher sensitivity to fixed-carbon starvation than *atg5-1* and *atg7-2* mutants, with 0% survival rate in *atg8-9m* mutants after seven days in darkness compared with 62% in *atg5-1* and 81% in *atg7-2* (Figures 2H and 2I). This higher sensitivity for *atg8-9m* was also demonstrated by a transient carbon starvation assay, in which light-grown seven-day-old seedlings were transferred to 1/2 MS medium without sucrose and kept in darkness (Jia *et al*., 2019). Compared with the WT and *atg7-2*, *atg8-9m* seedlings showed greatly accelerated senescence upon carbon deprivation (Figures 2J and 2K). Taken together, our data strongly suggest that the nine ATG8 isoforms are functionally redundant in plant autophagy, and the autophagic activity is completely dampened in the *atg8-9m* mutant.

### *atg8* Nonuple Mutant Showed Substantially Altered Proteome after Nitrogen and Fixed-Carbon Starvations

To thoroughly describe how the *Arabidopsis* proteome is affected by the *atg8* mutation and to relate it to the observed phenotypic changes, we analyzed the protein abundances in bulk by shotgun LC-MS/MS followed by label-free quantification based on precursor ion intensities in the first mass spectrometry dimension (MS1) (McLoughlin *et al*., 2018). In total, our quantitative MS data were collected for 5928 and 4322 proteins in N– and C-starved samples, respectively (Supplemental Table 2-4). Principal component analysis (PCA) showed that, consistent with previous *Arabidopsis* and maize proteomic studies (Avin-Wittenberg *et al*., 2015; McLoughlin *et al*., 2018; Havé *et al*., 2019), both genotype and nutrient starvation profoundly affected the *Arabidopsis* proteome profiles, with *atg8-9m* seedlings behaving similarly to *atg5-1* and *atg7-2* mutants in both non-starved and starved samples (Supplemental Figure 7). To better define how the *Arabidopsis* proteome was altered by *atg8* mutation and/or nutrient starvations, we generated volcano plots to analyze the behavior of all detected proteins of *atg8-9m* against wild-type plants (volcano plots analyzing of the proteomic profiles of *atg5-1* and *atg7-2* are shown in Supplemental Figure 7). As shown in Figure 3A and Supplemental Figure 7, whereas the total detected proteins were relatively evenly distributed between the wild type and *atg8-9m* in non-starved samples, the protein distribution was remarkably skewed toward the *atg8-9m* ledger under starved conditions. 17.3% (1,030) and 34.9% (1,511) of the detected proteins were significantly more abundant in N– and C-starved *atg8-9m* samples, respectively (P < 0.05; FDR < 0.05), whereas only 14.4% (854) and 9.5% (414) were significantly less abundant (Figure 3A). A similar skew of the proteome profile was also observed in starved *atg5-1* and *atg7-2* samples (Supplemental Figure 7). The levels of several proteins increased substantially in starved *atg8-9m* samples, especially under fixed-carbon conditions, such as the autophagy cargo receptor NBR1, and several complex/organellar proteins, including PAG1, PAE1, and RPT2a from the proteasome, ICL and PEX7 from the peroxisome, and MEE4 and TIM13 from the mitochondrion (Figure 3A).

The global effect of the *atg8-9m* mutation on organelle/complex profiles was further highlighted by GO term enrichments and volcano plots based on cellular compartment (Figure 3B and Supplemental Figure 8). Protein groups associated with plastid, ribosome, peroxisome, mitochondrion, ER, plasma membrane and proteasome complex were more abundant in all nutrient-starved *atg* mutants, although less prominent in N-starved samples. We also found that the chloroplast-associated proteins were less abundant in non-starved *atg8-9m* plants compared to wild-type ones. In addition, under fixed-carbon starvation conditions, the total detected proteins and proteins associated with several organelles, including ribosome, mitochondrion and plasma membrane were more abundant in *atg8-9m* compared with those of *atg5-1* and *atg7-2* mutants, which probably reflects a more sensitive phenotype of the *atg8-9m* mutant to fixed-carbon starvation (Figure 3A and Supplemental Figure 8). To further analyze the specific effects of the *atg8* mutation on compartments/processes within the *Arabidopsis* proteome, we performed a GO enrichment analysis of the proteins that were affected by the *atg8-9m* mutation but not paralleled by *atg5-1* or *atg7-2* (14.3% of total proteins analyzed in +/–N samples and 18.7% in +/–C samples). As shown in Figure 3C, a number of new GO terms emerged that became prominent in the *atg8-9m* mutants. Particularly notable were proteins associated with the trans-Golgi network and the Golgi apparatus regardless of nutrient stress (Figure 3D), consistent with a non-canonical role for ATG8 in maintaining Golgi hemostasis (Zhou *et al*., 2023).

### RABG3 Proteins Interact with ATG8 via the AIM-LDS Interface

To better understand the role of *Arabidospis* ATG8s in autophagosome maturation, we further investigated the ATG8-interacting partners involved in autophagosome-vacuole fusion. It has been reported that RABG3f, a member of the RAB7 GTPase family, was identified as an ATG8-interaction protein by yeast two-hybrid screening (Marshall *et al*., 2019), and ATG8 was revealed in the immunoprecipitants of YFP-RABG3f (Rodriguez-Furlan *et al*., 2019). Studies with the counterparts of RABG3f in yeast (Ypt7) and metazoans (RAB7) revealed that RAB7/Ypt7 is an important regulator of multiple endocytic and autophagic processes, including trafficking, maturation, lysosomal biogenesis and maintenance, and fusion (Zhao *et al*., 2021). Therefore, we first tested whether RABG3f specifically interacts with ATG8a *in planta* using an established luciferase complementation imaging (LCI) assay (Chen et al., 2008). We generated the constructs expressing RABG3f fused C-terminally to the N-terminal fragment of the LUC reporter (RABG3f-nLUC) and ATG8a fused N-terminally to the C-terminal fragment of the LUC reporter (cLUC-ATG8a). Empty nLUC and cLUC vectors were used as negative controls in this assay. When RABG3f-nLUC, cLUC-ATG8a and the control vectors were co-expressed in *N. benthamiana* leaves mediated via *A. tumefaciens*, only samples expressing the combination of RABG3f-nLUC and cLUC-ATG8a showed strong interaction as measured by bioluminescence signals, whereas samples expressing RABG3f-nLUC or cLUC-ATG8a with control vectors did not (Figure 4A). *In vitro* pull-down assay also confirmed the interaction of RABG3f with ATG8 (Figure. 4B). Further LCI assays showed that such interaction was conserved between these two protein families, as RABG3f could interact with the representative members from other major clade/subclade of the ATG8 family (ATG8e and ATG8h), and ATG8a could interact with other representative members of RABG3 family, including RABG3b, RABG3d and RABG3e (Supplemental Figure 9). In addition, we tested the interaction of ATG8a with RABG3f with different nucleotide binding states. When wild-type, GTP-bound, constitutive active RABG3f(CA) and GDP-bound, dominant-negative RABG3f(DN) variants were co-expressed together with ATG8, they all showed strong LUC complementation signals (Supplemental Figure 10A), suggesting that the interaction between ATG8a and RABG3f is independent of the activation of RABG3f. The expressions of RABG3/RABG3(CA)-nLUC, RABG3(DN)-nLUC, and cLUC-ATG8a were also validated by immunoblotting with anti-Luc (Supplemental Figure 10B).

**Figure 4.**
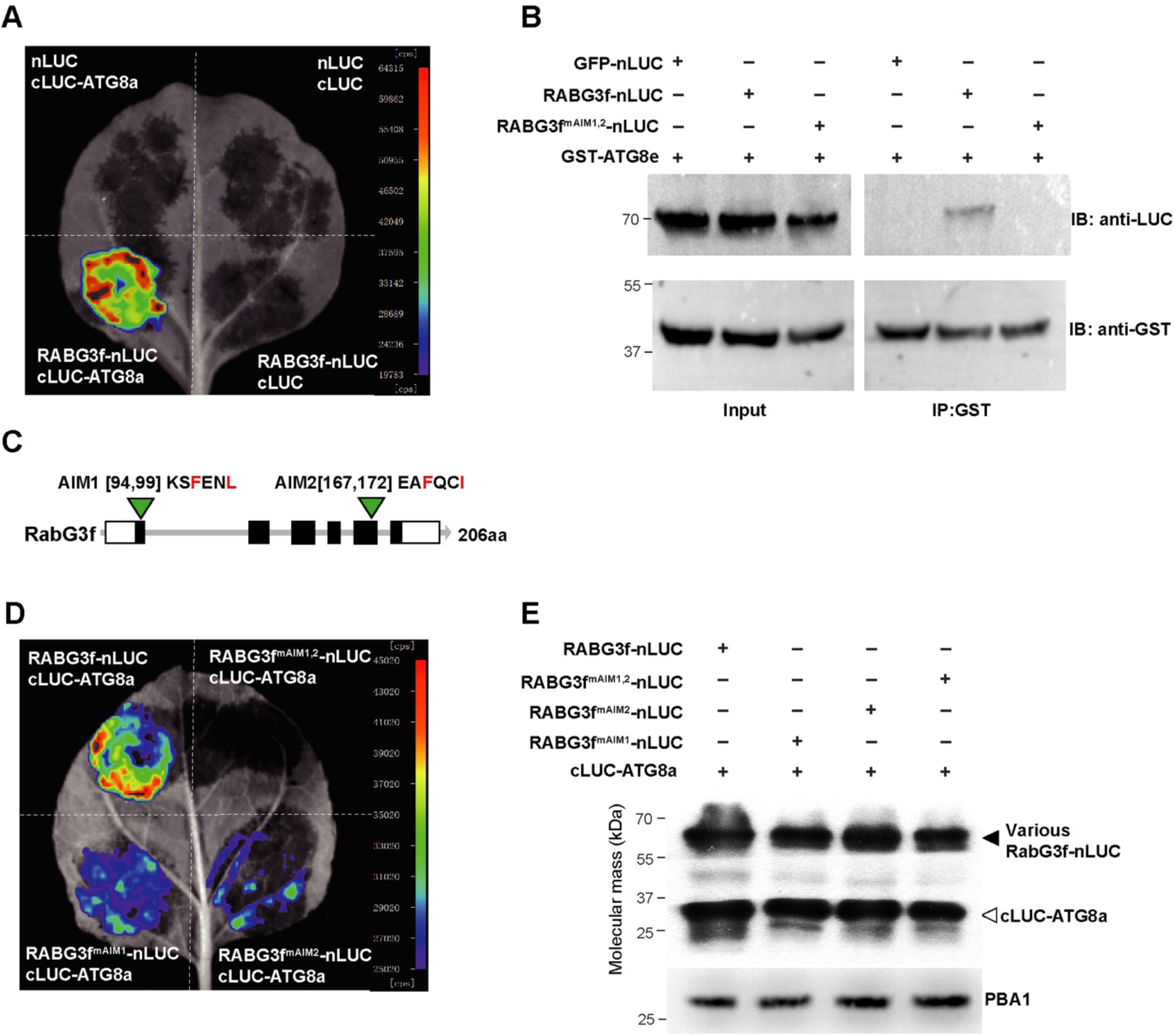
RABG3f Interacts with ATG8 Through Its ATG8-Interacting Motif (AIM). **(A)** Luciferase complementation imaging (LCI) assay to detect the interaction between RABG3f and ATG8a. nLUC and cLUC were used as negative controls. **(B)** *In vitro* pull-down assay. Protein extracts obtained from tobacco leaves infiltrated with the indicated constructs were incubated with the recombinant GST-ATG8e proteins coupled with glutathione beads to analyze the interaction between ATG8 and RABG3f. Both the input and pull-down samples were subjected to Western blotting using anti-luciferase or anti-GST. **(C)** Structure of the *RABG3f* gene including the ATG8-interacting motifs (AIMs) and the corresponding mutations in the AIMs. **(D)** Luciferase complementation between RABG3f and ATG8a was diminished by the AIM mutations in RABG3f. **(F)** The expression levels of cLUC and nLUC fusion proteins in (D) were validated by immunoblotting with anti-luciferase antibody, which recognizes both nLUC and cLUC of luciferase. Antibody against PBA1 was used as a loading control.

To dissect the residues of RABG3f that bind to ATG8a, we first performed iLIR analysis (Kalvari *et al*., 2014; https://ilir.warwick.ac.uk/index.php) and identified two potential AIMs that are well conserved in the RABG GTPase family (Supplemental Figure 11), named as AIM1 (KSFENL, amino acid [aa] 94-99 in RABG3f) and AIM2 (EAFQCI, aa 167-172 in RABG3f), respectively. To test the role of the AIMs in ATG8 binding, we mutated the key residues of AIM1 (RABG3f^mAIM1^: F96A L99A) or AIM2 (RABG3f^mAIM2^: F169A I172A) to alanine within full-length RABG3f (Figure. 4C). The LCI assay result showed that the mutation of AIM1 or AIM2 sequences individually decreased, and the mutation of both AIMs completely abolished the interaction between RABG3f and ATG8a (Figure 4D and 4E), indicating that both AIMs are required for the interaction. All the RABG3f and its AIM mutated variants were similarly stable when transiently expressed driven by the 35S promoter in *N. benthamiana* (Figure 4E). Thus, it is unlikely that differences in interaction with ATG8a are due to differences in gene expression or protein stability.

### RABG3f Localizes to Autophagosomal Membrane

Given that RABG3f interacts with ATG8a through an AIM-LDS interface, we hypothesized that RABG3f might be recruited by ATG8s to autophagosomes for fusion with the vacuole. To test this above hypothesis, we attempted to visualize the colocalization of mCherry-RABG3f with the autophagic marker GFP-ATG8a. As shown in Figure 4, ∼65.7% of mCherry-RABG3f puncta colocalized with GFP-ATG8a in N-starved root tip cells (Figures 5A and 5D). Remarkably, some of these mCherry-RABG3f labeled structures are ring-shaped (Figures 5B and 5C). However, after combined N-starvation and ConA treatment, the colocalization ratio between GFP-ATG8a and mCherry-RABG3f was significantly reduced to ∼4.2% (ratio of GFP-ATG8a colocalized mCherry-RABG3f puncta/total GFP-ATG8a puncta; Figure 5D), as most of the GFP-ATG8-labeled autophagic bodies inside the vacuole did not colocalize with the mCherry signal.

**Figure 5.**
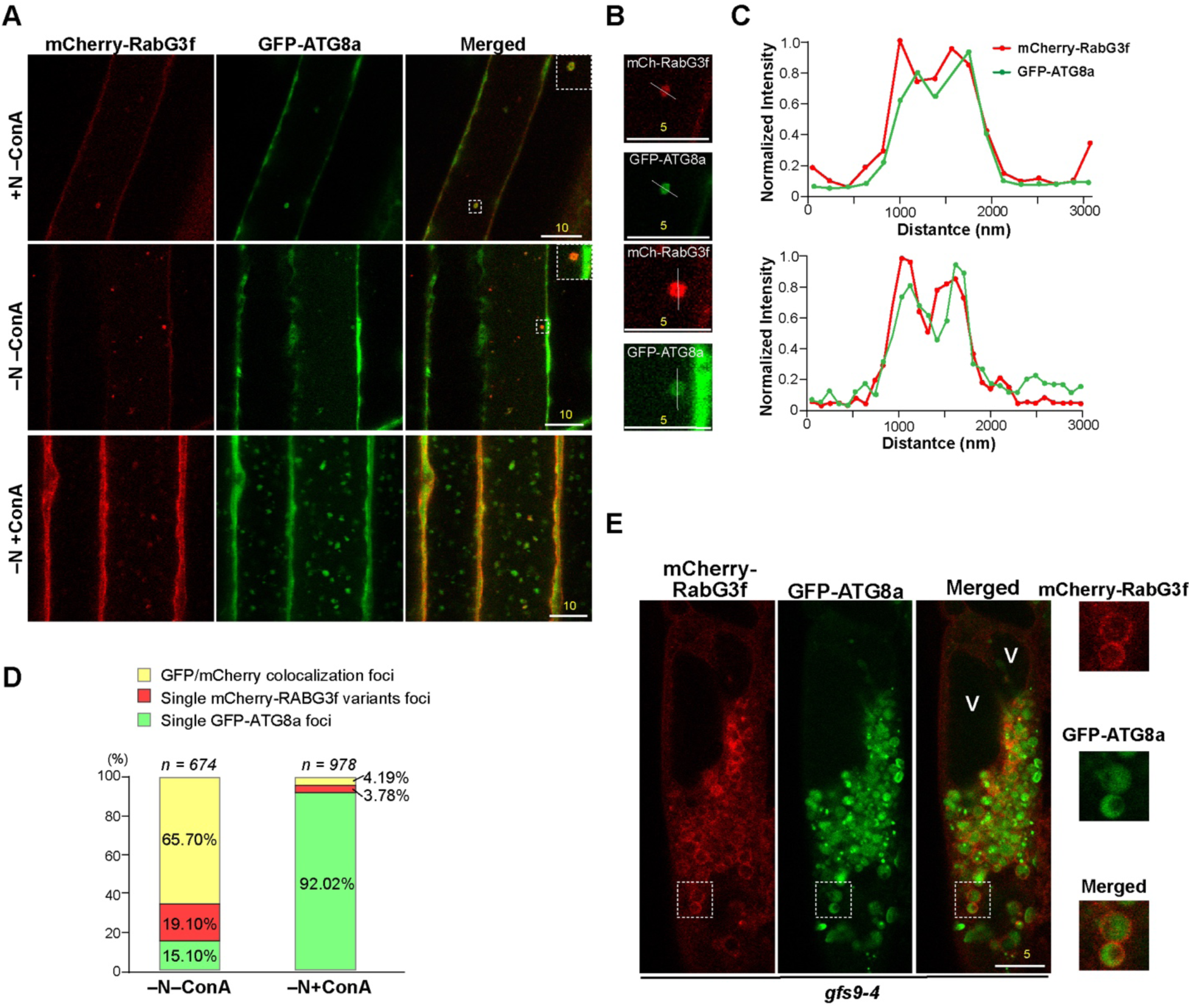
RABG3f Associates with Autophagic Vesicles. **(A)** mCherry-RABG3f colocalizes with GFP-ATG8a in autophagic vesicles. Wild-type plants stably expressing both reporters were grown on MS medium for 6 d and then transferred to nitrogen-deficient medium plus 1 μM ConA (–N+ConA) or DMSO (–N–ConA) for 8 h before confocal microscopy. Insets show 3x magnifications of the dashed boxes. Bars = 10 μm **(B-C)** Enlarged split-channel images corresponding to the dashed boxes in (A). Intensity profiles along the indicated white lines are plotted in (C). Bars = 5 μm **(D)** Quantification of the colocalization ratio between RABG3f and autophagosomal marker ATG8. A total of *n* = 674 (–N–ConA) and *n* = 978 (–N+ConA) particles were used for colocalization ratio calculations. **(E)** mCherry-RABG3f forms ring-like structures and colocalizes with GFP-ATG8a in the *gfs9-4* mutant. Mutant seedlings stably expressing both reporters were grown on MS medium for 6 d and then exposed to nitrogen-deficient medium containing 1 μM ConA for 8 h before confocal microscopy. The far right panels show the magnifications of the white dashed boxes. V: vacuole.

**Figure 6.**
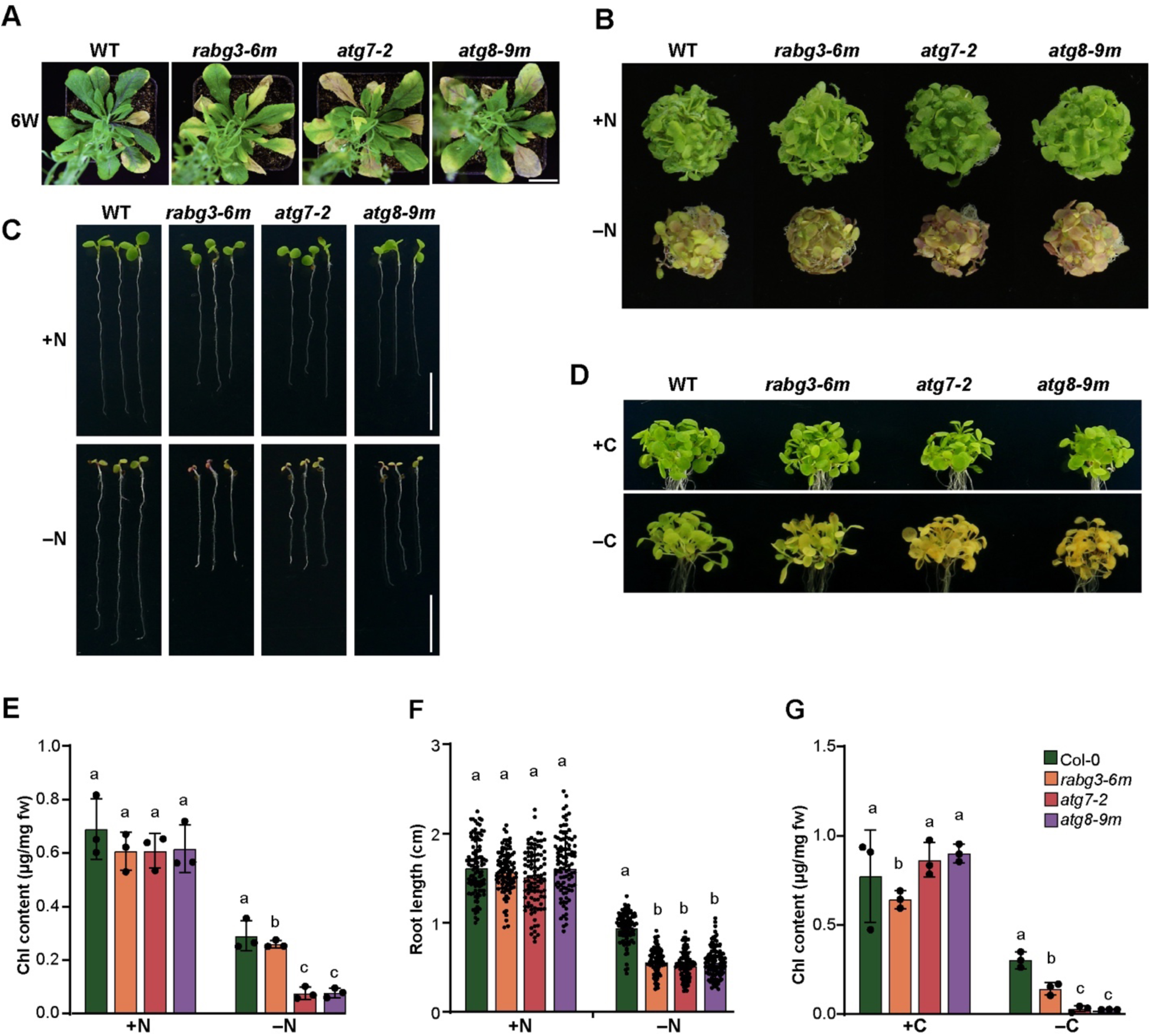
Phenotypic Analyses of the *Arabidopsis rabg3a,b,c,d,e,f* (*rabg3-6m*) Sextuple Mutant. The homozygous *rabg3-6m* mutant, the autophagy mutants *atg7-2* and *atg8-9m*, and WT were included for comparison. **(A)** Accelerated senescence. Plants were grown at 22°C in nutrient-rich soil under LD conditions for six weeks. Bar = 1 cm. **(B)** Enhanced sensitivity to nitrogen starvation. One-week-old seedlings were treated and observed as in Figure 2B. **(C)** Short primary root phenotype of *rabg3-6m* in response to nitrogen starvation. Seeds were grown as in Figure 2D. Bars = 5 mm. **(D)** Senescence phenotypes of the *rabg3-6m* mutant after dark treatment. Plants were grown and treated as in Figure 2J. **(E)** Total chlorophyll content of the plants shown in (B). Data are presented as mean ± s.d. (*n* = 3 biological replicates, 80-120 seedlings per genotype in each independent experiment). **(F)** Quantification of root length of plants shown in (C). Data are presented as mean ± s.d. (*n* = 3 biological replicates, 30-40 seedlings per genotype in each independent experiment). **(G)** Total chlorophyll content of the plants shown in (D). Data are presented as mean ± s.d. (*n* = 3 biological replicates, 80-120 seedlings per genotype in each independent experiment). Different letters in (E), (F), and (G) indicate significant differences (*P* < 0.05) as determined using two-way ANOVA followed by Tukey’s multiple comparison test.

To better visualize the subcellular localization of RABG3f in relation to the autophagosomes, we introduced both reporters into the *gfs9-4* mutant. Previous studies have shown that GFS9 is a peripheral protein of the Golgi apparatus, and its deficiency causes several membrane trafficking defects, including the abnormal accumulation of enlarged vesicles such as late endosomes and autophagosome-like structures (Ichino *et al*., 2014). When the autophagic marker GFP-ATG8a was introduced into the mutant, enlarged GFP-ATG8a dots were found to aggregate around the peri-vacuolar regions of *gfs9-4* root cells, in contrast to the smaller, vacuolar-resident dots observed in the wild type after nitrogen starvation and ConA treatment (Supplemental Figure 12). Under similar conditions, GFP-ATG8a– and mCherry-RABG3-positive structures were observed by confocal microscopy in the *gfs9-4* mutant, and mCherry-RABG3f signals formed ring-shaped structures surrounding the ATG8a-positive dots, suggesting that the RABG3f protein associates with the membrane of autophagosomes (Figure 5E).

### Loss of RABG3 GTPases Confers Hypersensitivity to Nutrient Stress

To determine what function RABG3 proteins might have in plant autophagy, we then challenged the sextuple mutant *rabg3a,b,c,d,e,f* (abbreviated as *rabg3-6m*) with a series of phenotypic analyses (Ebine *et al*., 2014). The autophagy-defective mutants *atg7-2* and *atg8-9m* were served as positive controls in these assays. When grown in nutrient-rich soil, *rabg3-6m* plants senesced earlier than wild type but slightly later than *atg7-2* and *atg8-9m* controls under LD conditions (Figure 6A). Such early senescence phenotypes become more pronounced when grown under SD conditions. Yellowing leaves were observed in ∼ten-week-old *rabg3-6m*, *atg7-2*, and *atg8-9m* mutants, whereas wild-type plants remained relatively green and healthy (Supplemental Figure 13).

When grown in nitrogen-limited MS medium, the *rabg3-6m* mutant also exhibited enhanced senescence-like symptoms compared with the wild type, and its chlorophyll content declined significantly faster than that of the wild type but slower than those of the *atg7-2* and *atg8-9m* mutants. (Figures 6B and 6E). Notably, the *rabg3-6m* mutant produced much shorter roots than wild-type plants when grown on nitrogen-depleted medium, resembling the *atg7-2* and *atg8-9m* mutants. In addition, a transient carbon starvation experiment was performed accordingly to test the hypersensitivity of *rabg3-6m* plants to fixed-carbon stress (Jia *et al*., 2019; Sun *et al*., 2023). Plants were grown on 1/2 MS medium supplemented with Suc under an LD photoperiod for one week, and then transferred to 1/2 MS medium without added Suc and kept in the dark for ten days. Compared with WT seedlings, *rabg3-6m* mutant seedlings exhibited a pronounced yellowing phenotype within ten days (Figures 6D and 6G). Taken together, loss of RABG3 proteins resulted in early senescence and increased sensitivity to nutrient stress.

### Autophagic Activity Is Downregulated in *rabg3-6m* Mutant

Since the *rabg3-6m* mutant is hypersensitive to nutrient (carbon and nitrogen) starvation, we asked whether this hypersensitivity to starvation is caused by a defect in autophagy. Therefore, GFP-ATG8a was transiently expressed in the leaf protoplasts prepared from wild-type and *rabg3-6m* plants (Figure 7A). GFP-ATG8a-labeled puncta in the vacuole of *rabg3-6m* cells were remarkably fewer (∼3.5-fold fewer) than in wild type upon ConA treatment (Figure 7B). Previous studies have shown that the lytic vacuole marker Aleu-GFP is trafficked to the central vacuole via the ER, Golgi and prevacuolar compartment (PVC), guided by a specific vacuolar sorting signal from the barley (*Hordeum vulgare*) protease Aleurain (Di Sansebastiano *et al*., 2001). When Aleu-GFP was transiently expressed in protoplasts, most WT cells showed a diffuse vacuolar fluorescence. In contrast, in *rabg3-6m* cells, bright punctate structures were detected in addition to the faint diffuse vacuolar fluorescence. (Supplemental Figure 14A). These punctate structures may represent Aleu-GFP abnormally accumulated in early endocytic compartments when its vacuolar trafficking is blocked, which has been observed in several *Arabidopsis* endocytic mutants (Sanmartin *et al*., 2007; Zeng *et al*., 2015; Kim *et al*., 2022). The block of Aleu-GFP transport in the *rabg3-6m* mutant was further validated using a GFP release immunoblot assay. As shown in Supplemental Figure 14B, while GFP was almost completely released in WT cells 8 h after transfection, a significant amount of Aleu-GFP was still detected in *rabg3-6m* cells. In contrast to the autophagic marker GFP-ATG8a and the vacuolar marker Aleu-GFP, the vacuolar trafficking of ABS3-GFP was not affected in *rabg3-6m* protoplasts, suggesting that the ABS3-ATG8 proteolysis pathway does not require the RABG3 GTPase (Supplemental Figure 14C and 14D).

**Figure 7.**
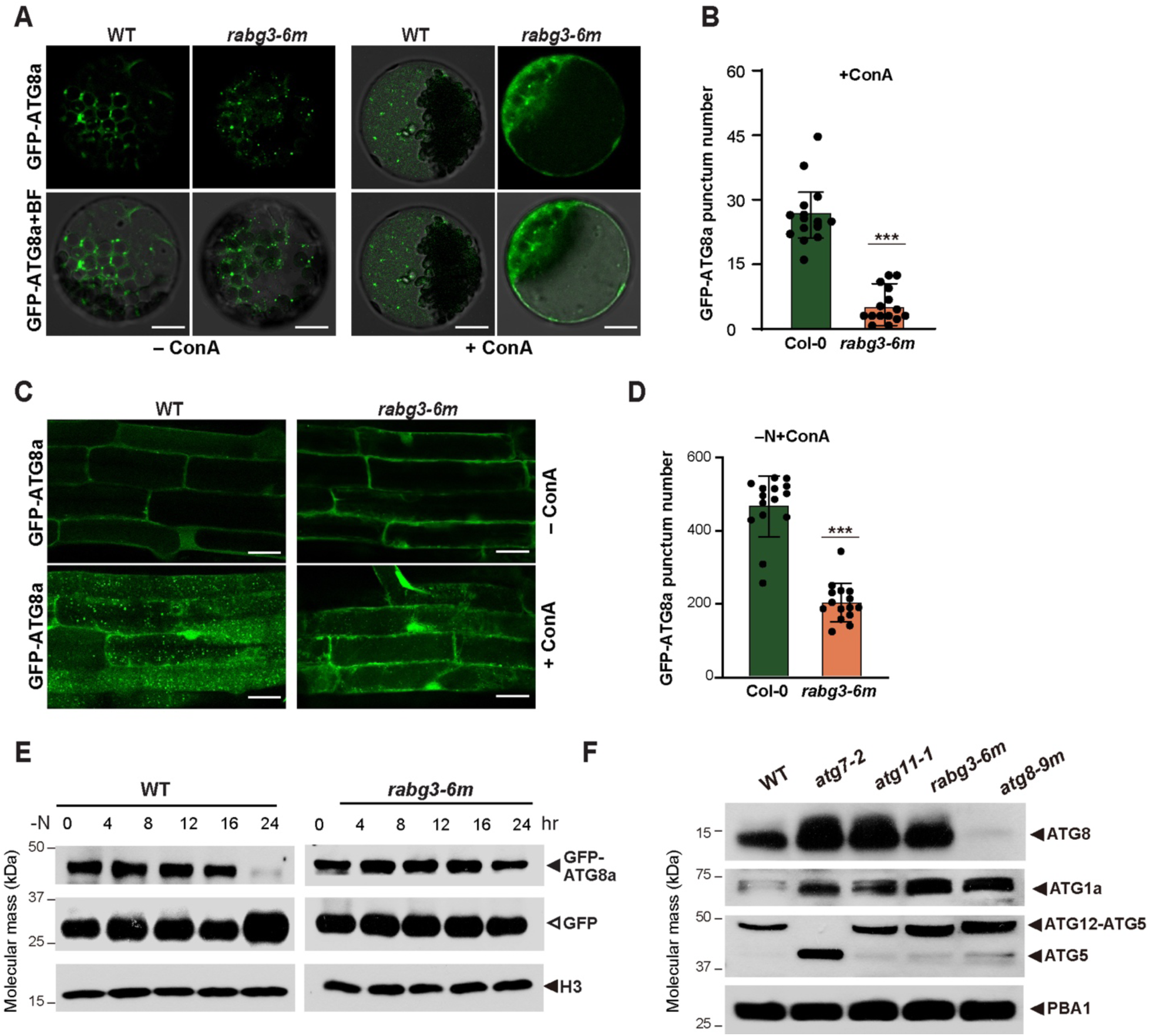
Lack of RABG3 Blocks Autophagic Body Deposition in the Vacuole. **(A-B)** Effects of *rabg3* mutations on the vacuolar transport of GFP-ATG8a in leaf protoplasts. Leaf protoplasts of WT and *rabg3-6m* mutant were transformed with the GFP-ATG8a construct and treated with 1 μM ConA for 12 to 14 h before confocal imaging analysis. Bars = 10 μm. Quantification of GFP-ATG8a puncta shown in (B). Images (*n* = 15) were collected to measure the number of puncta. Asterisked columns represent *rabg3-6m* mutants that are significantly different from WT, according to Student’s *t*-test. \*\*\**P* < 0.001. **(C-D)** Effects of *rabg3* mutations on the vacuolar deposition of GFP-ATG8a in stably transgenic plants. Seedlings expressing GFP-ATG8a were grown on nitrogen-containing MS solid medium for 6 d and then transferred to nitrogen-deficient medium supplemented with 1 μM ConA (– N+ConA) or DMSO (–N–ConA) for another 8 h before confocal fluorescence microscopy analysis of root cells. Quantification of GFP-ATG8a puncta shown in (D). Images (*n* = 15) were collected to measure the number of puncta. Asterisked columns represent *rabg3-6m* mutants that are significantly different from WT, according to Student’s *t*-test. \*\*\**P* < 0.001. **(E)** Immunoblot detection of the free GFP released during the vacuolar degradation of GFP-ATG8a. One-week-old wild-type and *rabg3-6m* seedlings described in (C) were transferred to fresh liquid MS medium without nitrogen (–N) for the indicated time. Total protein was subjected to immunoblot analysis using anti-GFP antibodies. The GFP-ATG8a fusion and free GFP are indicated by closed and open arrowheads, respectively. H3 was used to confirm approximately equal protein loading. **(F)** Immunoblot detection of multiple ATG proteins from wild-type (WT), *rabg3-6m*, *atg7-2*, and *atg8-9m* mutants using antibodies against ATG8, ATG1a and ATG5, respectively. Antibody against PBA1 was used as a loading control.

To further confirm this notion, we introduced GFP-ATG8a into the *rabg3-6m* background to monitor the autophagic flux by conventional Agrobacterium-mediated transformation. After combined nitrogen starvation and ConA treatment, GFP-ATG8a puncta were readily detected in WT root cells, whereas in *rabg3-6m*, vacuolar deposition of GFP-ATG8a puncta was inhibited; there were approximately 2.5-fold fewer GFP-ATG8a puncta in *rabg3-6m* compared with WT (Figure 7C and 7D). Consistent with this finding, the autophagic flux was apparently suppressed in the *rabg3-6m* mutant compared with WT, as evidenced by the decreased ratio of free GFP to GFP-ATG8a fusion, in response to nitrogen starvation (Figure 7E). In addition, we also measured the levels of different ATG proteins to determine where RABG3 affects the autophagy system. ATG8 and ATG1a proteins were significantly accumulated in the *rabg3-6m* mutant, consistent with previous observations with other autophagy mutants in which a defect in autophagy suppresses the turnover of these proteins (Thompson *et al*., 2005; Phillips *et al*., 2008; Chung *et al*., 2010; Li *et al*., 2014; Huang *et al*., 2019). In contrast, the conjugation of ATG12 to ATG5 is not affected in this mutant (Figure 7F). Taken together, these results suggest that loss of the RABG3 GTPase impair plant autophagy.

### RABG3-ATG8 Interaction Is Essential for the Vacuolar Deposition of Autophagic Vesicles

Given our discovery of the interaction of RABG3f with ATG8a and its association with the autophagosome, we ask whether the RABG3f-ATG8a interaction is essential for the vacuolar deposition of autophagosomes. To test this possibility, we first transiently co-expressed GFP-ATG8a with either mCherry-RABG3f or mCherry-RABG3f^mAIM1,2^ (AIM-mutant) in protoplasts prepared from *rabg3-6m* leaves. As shown in Figure 8A and 8B, the accumulation of GFP-ATG8a puncta in the vacuole was readily observed in the cells expressing mCherry-RABG3f after ConA treatment, whereas much fewer GFP-ATG8a puncta (∼12-fold fewer) were detected in similarly treated protoplasts expressing the mCherry-RABG3f^mAIM1,2^ variant, indicating that expression of mCherry-RABG3f, but not mCherry-RABG3f^mAIM1,2^, reversed the effect of the *rabg3-6m* mutation on the vacuolar deposition of autophagosomes. In contrast, when the vacuolar marker Aleu-GFP was coexpressed with either mCherry-RABG3f or mCherry-RABG3f^mAIM1,2^ variants, most *rabg3-6m* protoplasts expressing both constructs showed a strong diffuse green vacuolar fluorescence regardless of the RABG3f variants expressed (Figure 8D). Consistently, the inhibition of GFP processed from the Aleu-GFP fusion in the *rabg3-6m* mutant could be reversed by expressing either mCherry-RABG3f or mCherry-RABG3f^mAIM1,2^(Figure 8E), suggesting that the endocytic trafficking of Aleu-GFP is independent of the RABG3f-ATG8 interaction.

**Figure 8.**
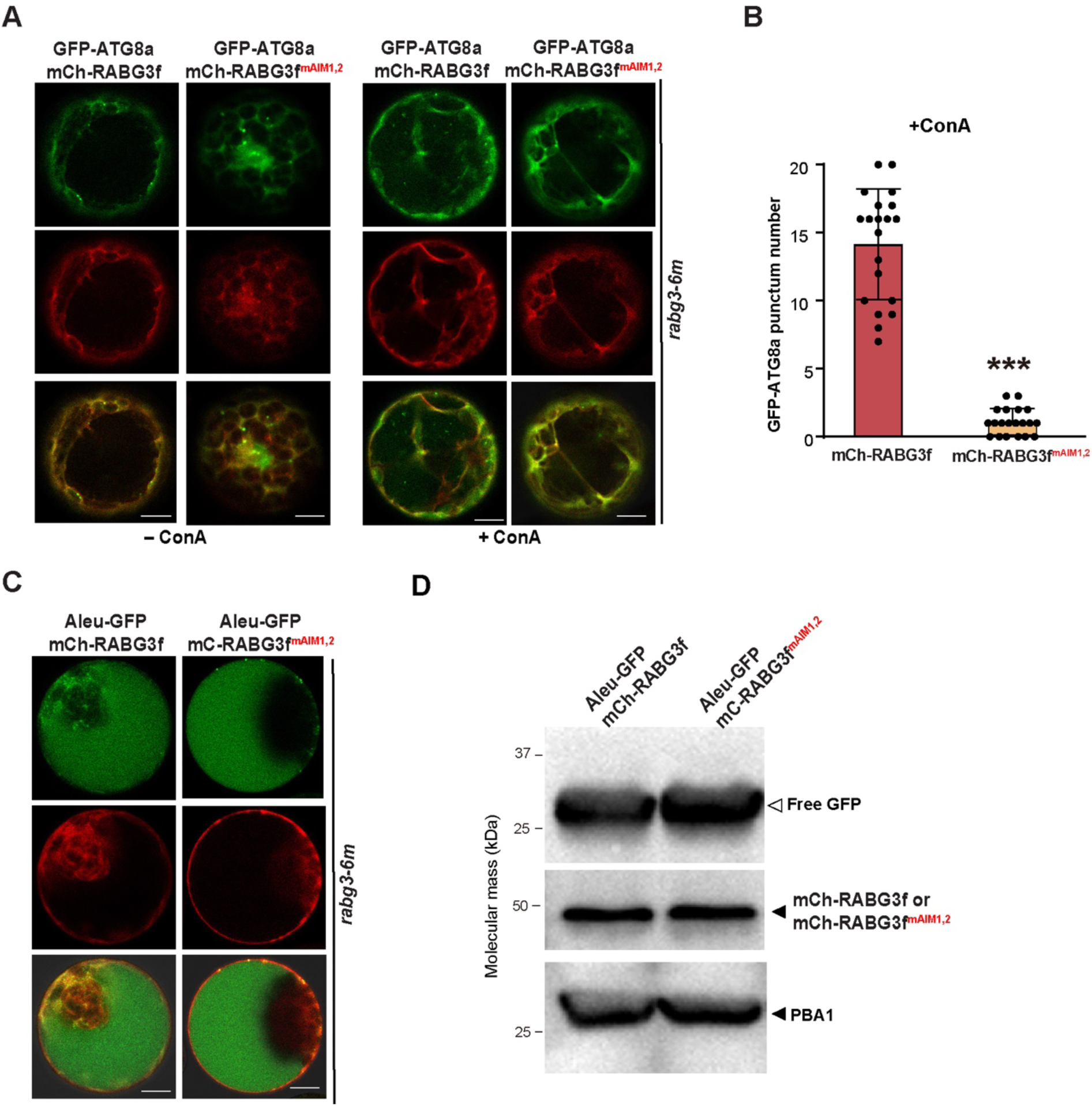
RABG3f Mediates the Vacuolar Deposition of GFP-ATG8a via Its AIMs. **(A-B)** Confocal images of protoplasts prepared from *rabg3-6m* transiently coexpressing GFP-ATG8a with either mCherry-RABG3f or mCherry-RABG3f^mAIM1,2^. Leaf protoplasts were incubated in liquid medium containing DMSO (–ConA) or ConA (B) for 12 h prior to observation. Bars = 10 μm. Quantification of vacuolar puncta of ABS3-GFP shown in (B). Images (*n* = 20) were collected to measure the number of puncta. Asterisked columns represent protoplasts expressing the mCherry-RABG3f^mAIM1,2^ construct that are significantly different from those expressing the wild-type form of RABG3f, according to Student’s *t*-test. \*\*\**P* < 0.001. **(C)** Confocal images of *rabg3-6m* protoplasts transiently expressing Aleu-GFP with either mCherry-RABG3f or mCherry-RABG3f^mAIM1,2^. Leaf protoplasts were incubated in liquid medium for 8 h prior to observation. Bars = 10 μm. **(E)** Immunoblot analysis of *Arabidopsis* leaf protoplasts transiently expressing Aleu-GFP with either mCherry-RABG3f or mCherry-RABG3f^mAIM1,2^ as shown in (D) using anti-GFP and anti-mCherry antibodies. PBA1 was used to confirm approximately equal protein loading.

To further investigate whether the RABG3-ATG8 interaction is essential for the autophagic pathway in *Arabidopsis,* and to confirm the *rabg3-6m* phenotypes by complementation, we constructed two complementation lines. One was performed with wild-type RABG3f driven by the *Arabidopsis* UBQ10 promoter, while the other was complemented with the AIM-mutated RABG3f^mAIM1,2^ variant (Supplemental Figure 15A). Immunoblot analysis revealed that mCherry-RABG3f or mchery-RABG3f^mAIM1,2^ was coexpressed with the autophagic marker GFP-ATG8a in the resulting homozygous plants (Supplemental Figure 15B). Phenotyping analyses revealed that the complemented line with wild-type RABG3f exhibited similar to the WT in all tested conditions, including senescence, nitrogen-limiting, and carbon-limiting conditions. However, the complemented line with the AIM-mutated RABG3f ^mAIM1,2^ variant still behaved like the *rabg3-6m* mutant (Figure 9A-G). Interestingly, this reporter successfully rescued the defects in the trafficking and processing of 12S globulin precursor in the *rabg3-6m* mutant (Supplemental Figure 15C), which is consistent with the observation of the trafficking and processing of Aleu-GFP, and further suggests that this RABG3f^mAIM1,2^ variant can rescue the endocytosis function of *Arabidopsis* RAB7/RABG3s. We further investigated whether there was a difference in the autophagy pathway between the complementation lines. As shown in Figure 9I-J, confocal imaging analysis and quantification revealed that most mCherry-RABG3f reporter (74.32%) colocalized with GFP-ATG8a under nitrogen-limiting conditions, whereas only 5.71% colocalization foci of both reporters were found in the RABG3f^mAIM1,2^ complementation line, further confirming the importance of RABG3-ATG8 interaction for RABG3 association with autophagic vesicles.

**Figure 9.**
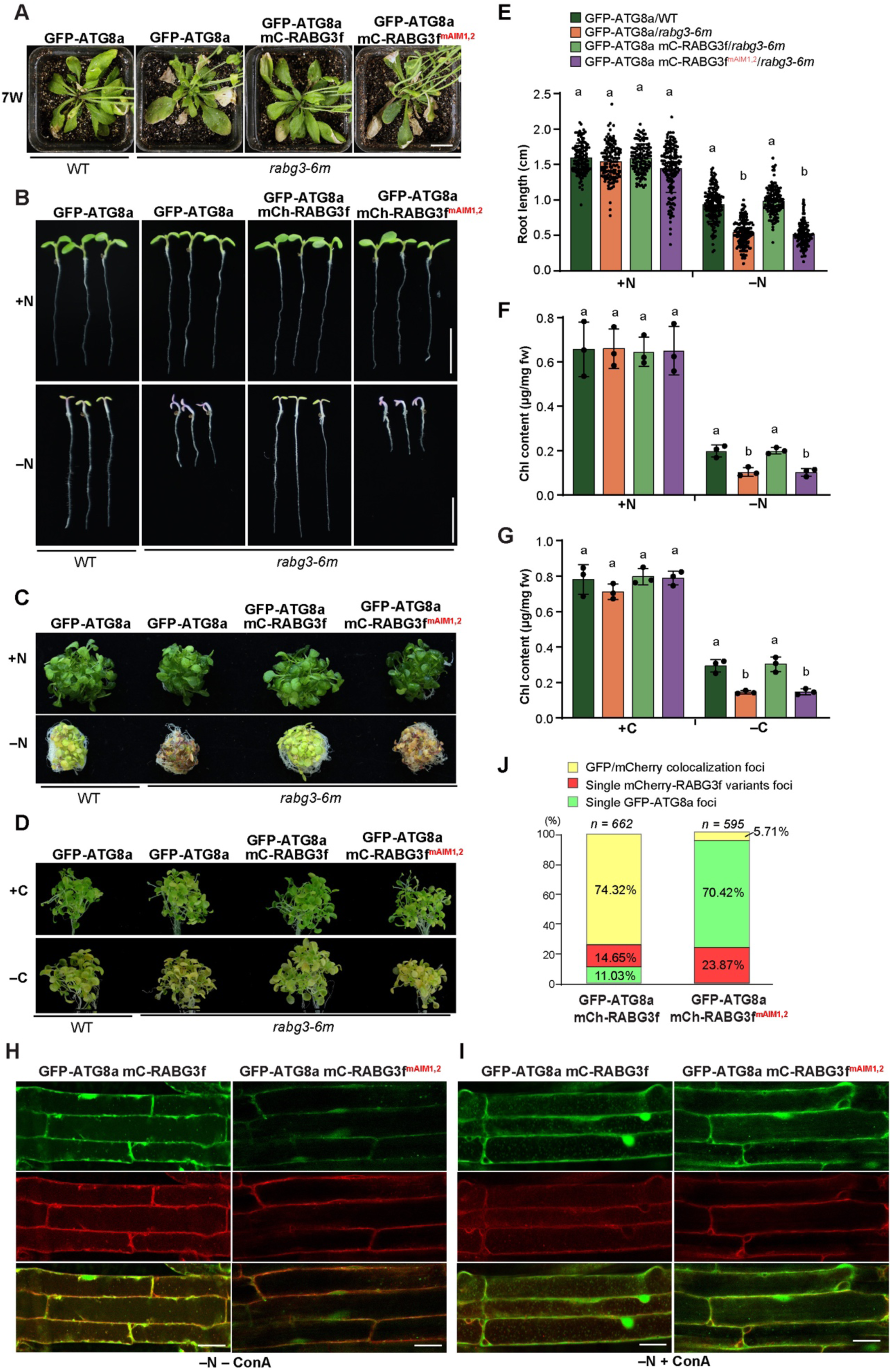
Complementation Analysis of Wild-Type and AIM-Mutated RABG3f. (**A-G**) Wild-type but not AIM-mutated RABG3f rescues the early senescence and nutrient hypersensitivity of *rabg3-6m*. Phenotypic analyses were conducted as described in Figure 2. **(H-J)** Confocal fluorescence images of *rabg3-6m* root cells coexpressing GFP-ATG8a with either mCherry-RABG3f or mCherry-RABG3f^mAIM1,2^. Six-day-old seedlings were transferred to nitrogen-deficient medium supplemented with or without 1 μM ConA for another 12 h prior to confocal microscopic observation. Quantification of GFP-ATG8a puncta shown in (J). Images (*n* = 15) were collected to measure the number of puncta. Different letters in (E), (F), and (G) indicate significant differences (*P* < 0.05) as determined using two-way ANOVA followed by Tukey’s multiple comparison test.

## Discussion

Defining the role of ATG8s in plant autophagy has been difficult due to the genetic redundancy among the large number of *ATG8* genes in plants. To date, most of the knowledge about plant ATG8s has come from studies using mutants defective in ATG8 lipidation or plants overexpressing ATG8s. We have overcome this hurdle in *Arabidopsis* by systematically mutating the *ATG8* genes through CRISPR/Cas9-mediated mutagenesis (Figure 1 and Supplemental Figure 1). From the phenotypic analysis of the different *atg8* mutants, it appears that *Arabidopsis* ATG8s function redundantly in bulky autophagy, which is different from mammalian ATG8 members, which play distinct roles in the process of autophagosome biogenesis (Weidberg *et al*., 2010). The double, triple, quadruple, and quintuple mutants did not show obvious differences from WT plants in growth under all nutrient-starved conditions tested or in senescence (Figure 2). However, *Arabidopsis* plants in which all clade I *ATG8* genes were inactivated (*atg8* septuple) displayed phenotypes of hypersensitivity to nutrient starvation and early senescence, suggesting that the ATG8s function in plant autophagy in a gene-dosage dependent manner. The typical autophagy phenotypes of *atg8-7m* were consistent with the phenotypes of *Arabidopsis atg4ab* double mutants, in which the clade II isoforms ATG8h and ATG8i are synthesized with carboxyl-terminal Gly and theoretically do not require the activity of ATG4 (Yoshimoto *et al*., 2004; Chung *et al*., 2010). Notably, we found that *atg8-9m* are more sensitive to fixed-carbon starvation than those mutants that abolish ATG8/12 conjugation (e.g., *atg7-2* and *atg5-1*), suggesting additional roles for ATG8 in carbon starvation-induced senescence that do not involve ATG8 lipidation. A recent study reported that the ABS3 subfamily of MATE proteins promotes carbon starvation-induced senescence via a ATG8-ABS3 interaction and subsequent deposition of ABS3 in the vacuole which does not require the ATG8 conjugation machinery (Jia *et al*., 2019). However, it remains unclear whether the hypersensitivity to carbon starvation in *atg8-9m* is due to disrupted trafficking of ABS3. In the future, it will be of great interest to investigate the molecular mechanism of the non-canonical ATG8-ABS3 interaction that regulates carbon starvation-induced senescence.

In *Arabidopsis* and certain mammalian cells types, most of the ATG5 is conjugated to ATG12, and free ATG5 is barely detectable regardless of nutrient availability (Mizushima *et al*., 2001; Thompson *et al*., 2005). Moreover, the formation of the ATG12–ATG5 conjugate proceeds normally even in some mutants of the autophagy core pathway, including those affecting the ATG1/13 kinase complex, the ATG2/9/18 transmembrane complex, and the phosphatidylinositol 3-kinase (PI3K) complex (Huang *et al*., 2019; Liu *et al*., 2020). The assembly of the ATG12–ATG5 conjugate was found to be disturbed only in mutants defective in ATG7 and ATG10, which function as E1 and E2 enzymes for the conjugation, respectively (Thompson *et al*., 2005; Phillips *et al*., 2008 and Figure 1C). Our data showed that the free ATG5 pool increases in *atg8* nonuple mutants (Figure 1C and Figure 7F), suggesting that the activity of the conjugation machinery may decrease in this mutant or that there may be a negative feedback regulatory loop may exist on ATG8 conjugation.

Structurally, all ATG8s have ubiquitin-like cores with an N-terminal extension that form two deep hydrophobic pockets for docking the AIMs of various cargos/receptors (Johansen & Lamark, 2020). Therefore, the structural differences in the hydrophobic pockets and the N-termini of ATG8s determine their binding specificity for interacting proteins during selective autophagy. Similar to the metazoan ATG8s, recent studies have begun to reveal the functional specialization of plant ATG8s. Interactome studies and domain swap analysis have shown that a single amino acid polymorphism at the N-terminal β-strand of potato ATG8CL confers binding specificity to a pathogen effector protein (Zess *et al*., 2019). CLC2, a light chain subunit of the *Arabidopsis* clathrin complex has been reported to interact specifically with clade II ATG8 isoforms and to mediate the plant immunity against the fungal pathogen *Golovinomyces cichoracearum* (Lan *et al*., 2024). Using NBR1 as a representative autophagic receptor, we showed that its trafficking was impaired only in *atg8-7m* and *atg8-9m* protoplasts, but not in other *atg8* mutants (Figure 1D and 1F). This probably reflects the broad binding specificity of NBR1 AIMs, as reported previously (Svenning *et al*., 2011). Nevertheless, a series of *Arabidopsis* mutants lacking different combinations of ATG8s, generated in this study, now allows the study of their functional specialization. Combined with mutagenesis and protein-protein interaction analyses, it should now be possible to understand the precise roles of *Arabidopsis* ATG8s in autophagy and other cellular processes.

Given the critical role of RAB7 in autophagosome-lysosome/vacuole fusion, it is, therefore, interesting to ask how RAB7 and/or its GEF, the MON1-CCZ1 complex are recruited to autophagosomal membrane. A yeast study showed that the MON1-CCZ1 complex is targeted to the autophagosome through the interaction between CCZ1 and Atg8 and subsequently promotes RAB7/Ypt7 recruitment (Gao *et al*., 2018). In addition, a study of fruit fly (*Drosophila melanogaster*) adipocytes proposed that phosphatidylinositol 3-phosphate (PI3P), produced by the ATG14-containing PI3K complex, is required for the autophagosomal association of the MON1-CCZ1 complex and RAB7 (Hegedűs *et al*., 2016). A similar finding was also obtained in the tobacco study; ATG14, paired with UVRAG (ultraviolet resistance-associated gene), another subunit of the class III PI3K complexes, to regulate autophagosome maturation by recruiting RAB7 and HOPS (Wang *et al*., 2022). Recently, Zhang *et al*. (2024) reported that the rice MON1-CCZ1 complex is involved in autophagy, most likely through the direct interaction of ATG8 with MON1 rather than CCZ1. In addition, rice RAB7 was also found to interact directly with ATG8. Here, we identified *Arabidopsis* RABG3f as an interacting partner of ATG8 via the AIM-LDS interface, which is critical for targeting RABG3f to the autophagosome membrane. Interestingly, we found that the interaction of RABG3f with ATG8 is independent of its GTP-binding state in the LCI assay (Supplemental Figure 10), although this may not represent its physiologically natural state since both RABG3f and ATG8 were transiently overexpressed. Moreover, we cannot exclude that other factors, such as the MON1-CCZ1 complex and/or UVRAG, are required for the recruitment and/or activation of RAB7 to autophagosomes. Nevertheless, the precise mechanisms by which RAB7 is recruited and activated during plant autophagy remain an area of active investigation.

Due to genetic redundancy, constitutively active or dominant negative RABG3 variants have been used to study the role of RAB7 in plant autophagy. For example, it was reported that overexpression of the RabG3b(CA) variant in *Arabidopsis* induced the formation of autophagic vesicles in developing tracheary element cells (Kwon *et al*., 2010). In *N*. *benthamiana*, overexpression of the DN mutants of RABG3a or RABG3f impaired autophagic flux, as evidenced by the abnormal accumulation of NBR1 and autophagic vesicles, and reduced GFP-ATG8 processing (Wang *et al*., 2022). Consistent with these observations, our studies here showed that *rabg3-6m* exhibited prominent autophagy mutant phenotypes, including hypersensitivity to nitrogen and fixed-carbon starvations and premature rosette senescence (Figure 6). Autophagic flux was also subtly suppressed, as shown by the immunoblot analysis of ATG proteins, confocal microscopy of the GFP-ATG8a reporter, and the free GFP release assay (Figure 7). The relatively mild phenotypes observed in this mutant line could be due to residual expression of RABG3f or genetic redundancy of the other two family members, RABG1 and RABG2 (Ebine *et al*., 2014).

Further investigation of higher order RABG mutants may help to understand the contribution of RAB7 family proteins to the plant autophagic process.

In conclusion, in this study we have succeeded in generating the nonuple mutant line *atg8-9m* by combining mutations that disrupt both clade I and II ATG8s. Using the mutants lacking different ATG8s, we demonstrated that two clades of ATG8s redundantly control the autophagy activity in *Arabidopsis*. In addition, we uncovered the GTPase RABG3f as an interacting partner of ATG8 and the underlying mechanism for its targeting to the autophagosome. Further investigation of how ATG8, the MON1-CCZ1 complex, and other factors such as UVRAG and ATG14 act in concert to regulate the recruitment and activation of RAB7 to autophagosomes will lead to a better understanding of autophagosome-lysosome/vacuole fusion in plants.

## ACKNOWLEDGEMENTS

This work was supported by grants from the National Natural Science Foundation of China (grant 32370352 to F.Q.L. and grant 32070195 to X. H.), and the Natural Science Foundation of Guangdong Province (2024A1515011671) to X. H. We thank Drs. Hongbo Li, Cao Yang, and Caiji Gao (South Normal University, China) for technical support.

## AUTHOR CONTRIBUTIONS

F.Q.L., X. H., and R.D.V. designed the research. K.D., G.D., and Y.L. performed most of the experiments. K-E.C. performed the proteomic analysis. H.W., X.E. H., W.Y. H., and P.Z. provided technical support. T.U. provided the *rabg-6m* mutant and technical help to analyze data. F.Q.L., X. H., and R.D.V. wrote the manuscript with input from all authors.

## Supplemental Data

The following materials are available in the online version of this article.

**Supplemental Table 1.** Oligonucleotide Primers Used in The Study

**Supplemental Table 2.** Normalization Protein List and Values Used for Sample Correction.

**Supplemental Table 3.** Proteome Analysis.

**Supplemental Table 4.** Proteome Raw Data.

**Supplemental Table 5.** Statical analysis.

**Supplemental Figure 1.** Mutations in Various *atg8* Mutants.

**Supplemental Figure 2.** Transient Expression of Different GFP-Fused ATG Proteins in *atg8-9m* Protoplasts.

**Supplemental Figure 3.** Transient Expression of the GFP-NBR1 Fusion Protein in Protoplasts of Various *atg8* Mutants.

**Supplemental Figure 4.** Transient Expression of GFP-Fused ABS3 Protein in *atg8-9m* Protoplasts.

**Supplemental Figure 5.** Phenotypic Analysis of *Arabidopsis atg8* Mutants.

**Supplemental Figure 6.** *atg8* Mutants Produce Viable Pollen.

**Supplemental Figure 7**. Proteomic Comparisons of Wild Type and *atg* Mutants During N-or C-starvation.

**Supplemental Figure 8.** The Influence of *atg8* Mutation on Protein Abundances in Specific Cellular Compartments/Complexes in N-or C-starved *Arabidopsis* Samples.

**Supplemental Figure 9.** LCI Assays for the Detection of RABG3-ATG8 Interactions.

**Supplemental Figure 10.** Mutations in the GTPase Region of RABG3f Do Not Affect the RABG3f-ATG8a Interaction.

**Supplemental Figure 11.** Sequence Alignment of the *Arabidopsis* RABG Proteins.

**Supplemental Figure 12.** Intracellular Distribution of the Autophagy Marker GFP-ATG8a in the Root Elongation and Maturation Zones of *gfs9-4* Seedlings.

**Supplemental Figure 13.** The *rabg3-6m* Mutant Exhibited an Early Leaf Senescence Phenotype Under SD Conditions.

**Supplemental Figure 14.** RABG3 Is Required for the Vacuolar Deposition of Aleu-GFP But Not for ABS3-GFP.

**Supplemental Figure 15.** Genetic Complementation of *Arabidopsis rabg3-6m* with Wild-Type or AIM-Mutated RABG3f.

