## Supplemental Figures for "Genetic Analysis of *Arabidopsis* Autophagy-Related 8 Family Proteins Reveals Their Redundant and Regulatory Roles in Plant Autophagy"

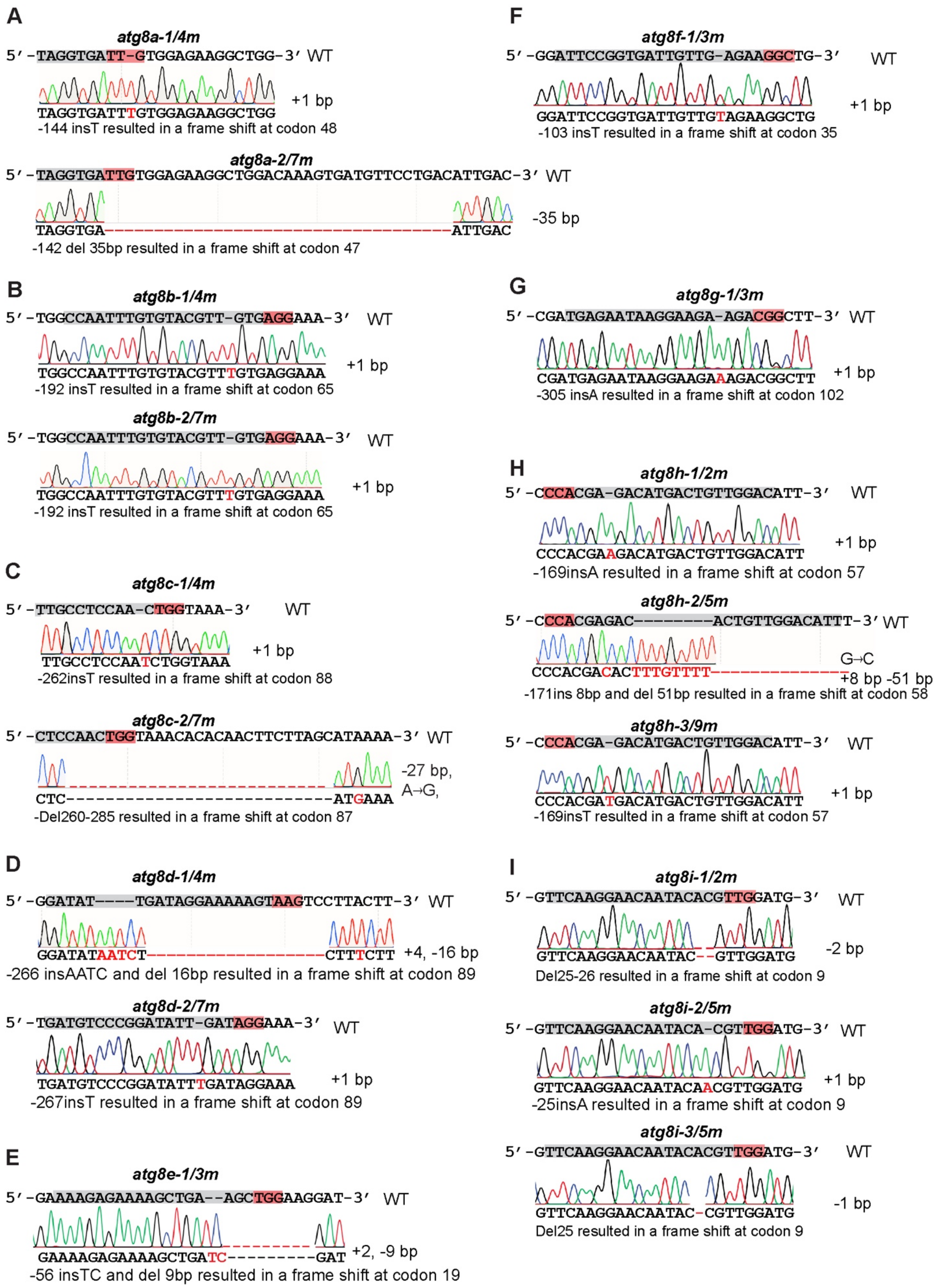
A
atg8a-1/4m
5′-TAGGTGATT-GTGGAGAAGGCTGG-3′ WT
+1 bp
TAGGTGATTTGTGGAGAAGGCTGG
-144 insT resulted in a frame shift at codon 48
atg8a-2/7m
5′-TAGGTGATTGTGGAGAAGGCTGGACAAAGTGATGTTCCTGACATTGAC-3′ WT
-35 bp
TAGGTGA-----------------------------------ATTGAC
-142 del 35bp resulted in a frame shift at codon 47
B
atg8b-1/4m
5′-TGGCCAATTTGTGTACGTT-GTGAGGAAA-3′ WT
+1 bp
TGGCCAATTTGTGTACGTTTGTGAGGAAA
-192 insT resulted in a frame shift at codon 65
atg8b-2/7m
5′-TGGCCAATTTGTGTACGTT-GTGAGGAAA-3′ WT
+1 bp
TGGCCAATTTGTGTACGTTTGTGAGGAAA
-192 insT resulted in a frame shift at codon 65
C
atg8c-1/4m
5′-TTGCCTCCAA-CTGGTAAA-3′ WT
+1 bp
TTGCCTCCAATCTGGTAAA
-262insT resulted in a frame shift at codon 88
atg8c-2/7m
5′-CTCCAACTGGTAAACACACAACTTCTTAGCATAAAA-3′ WT
-27 bp,
A→G,
CTC---------------------------ATGAAA
-Del260-285 resulted in a frame shift at codon 87
D
atg8d-1/4m
5′-GGATAT----TGATAGGAAAAAGTAAGTCCTTACTT-3′ WT
+4, -16 bp
GGATATAATCT----------------CTTTCTT
-266 insAATC and del 16bp resulted in a frame shift at codon 89
atg8d-2/7m
5′-TGATGTCCCGGATATT-GATAGGAAA-3′ WT
+1 bp
TGATGTCCCGGATATTTGATAGGAAA
-267insT resulted in a frame shift at codon 89
E
atg8e-1/3m
5′-GAAAAGAGAAAAGCTGA--AGCTGGAAGGAT-3′ WT
+2, -9 bp
GAAAAGAGAAAAGCTGATC---------GAT
-56 insTC and del 9bp resulted in a frame shift at codon 19
F
atg8f-1/3m
5′-GGATTCCGGTGATTGTTG-AGAAGGCTG-3′ WT
+1 bp
GGATTCCGGTGATTGTTGTAGAAGGCTG
-103 insT resulted in a frame shift at codon 35
G
atg8g-1/3m
5′-CGATGAGAATAAGGAAGA-AGACGGCTT-3′ WT
+1 bp
CGATGAGAATAAGGAAGAAAGACGGCTT
-305 insA resulted in a frame shift at codon 102
H
atg8h-1/2m
5′-CCCACGA-GACATGACTGTTGGACATT-3′ WT
+1 bp
CCCACGAAGACATGACTGTTGGACATT
-169insA resulted in a frame shift at codon 57
atg8h-2/5m
5′-CCCACGAGAC--------ACTGTTGGACATTT-3′ WT
G→C
+8 bp -51 bp
CCCACGACACTTTGTTTT--------------
-171ins 8bp and del 51bp resulted in a frame shift at codon 58
atg8h-3/9m
5′-CCCACGA-GACATGACTGTTGGACATT-3′ WT
+1 bp
CCCACGATGACATGACTGTTGGACATT
-169insT resulted in a frame shift at codon 57
I
atg8i-1/2m
5′-GTTCAAGGAACAATACACGTTGGATG-3′ WT
-2 bp
GTTCAAGGAACAATAC--GTTGGATG
Del25-26 resulted in a frame shift at codon 9
atg8i-2/5m
5′-GTTCAAGGAACAATACA-CGTTGGATG-3′ WT
+1 bp
GTTCAAGGAACAATACAACGTTGGATG
-25insA resulted in a frame shift at codon 9
atg8i-3/5m
5′-GTTCAAGGAACAATACACGTTGGATG-3′ WT
-1 bp
GTTCAAGGAACAATAC-CGTTGGATG
Del25 resulted in a frame shift at codon 9

**Supplemental Figure 1.** Mutations in Various *atg8* Mutants.

**(A)** Mutations in *ATG8a* of *atg8-4m* and *atg8-7m* mutants.

**(B)** Mutations in *ATG8b* of *atg8-4m* and *atg8-7m* mutants.

**(C)** Mutations in *ATG8c* of *atg8-4m* and *atg8-7m* mutants.

**(D)** Mutations in *ATG8d* of *atg8-4m* and *atg8-7m* mutants.

**(E)** Mutations in *ATG8e* of *atg8-3m* mutant.

**(F)** Mutations in *ATG8f* of *atg8-3m* mutant.

**(G)** Mutations in *ATG8g* of *atg8-3m* mutant.

**(H)** Mutations in *ATG8h* of *atg8-2m*, *atg8-5m*, and *atg8-9m* mutants.

**(I)** Mutations in *ATG8i* of *atg8-2m*, *atg8-5m*, and *atg8-9m* mutants. Red boxes: PAM sequences; grey boxes: ATG8-sgR target sequences; "+": insertion; "-": deletion.

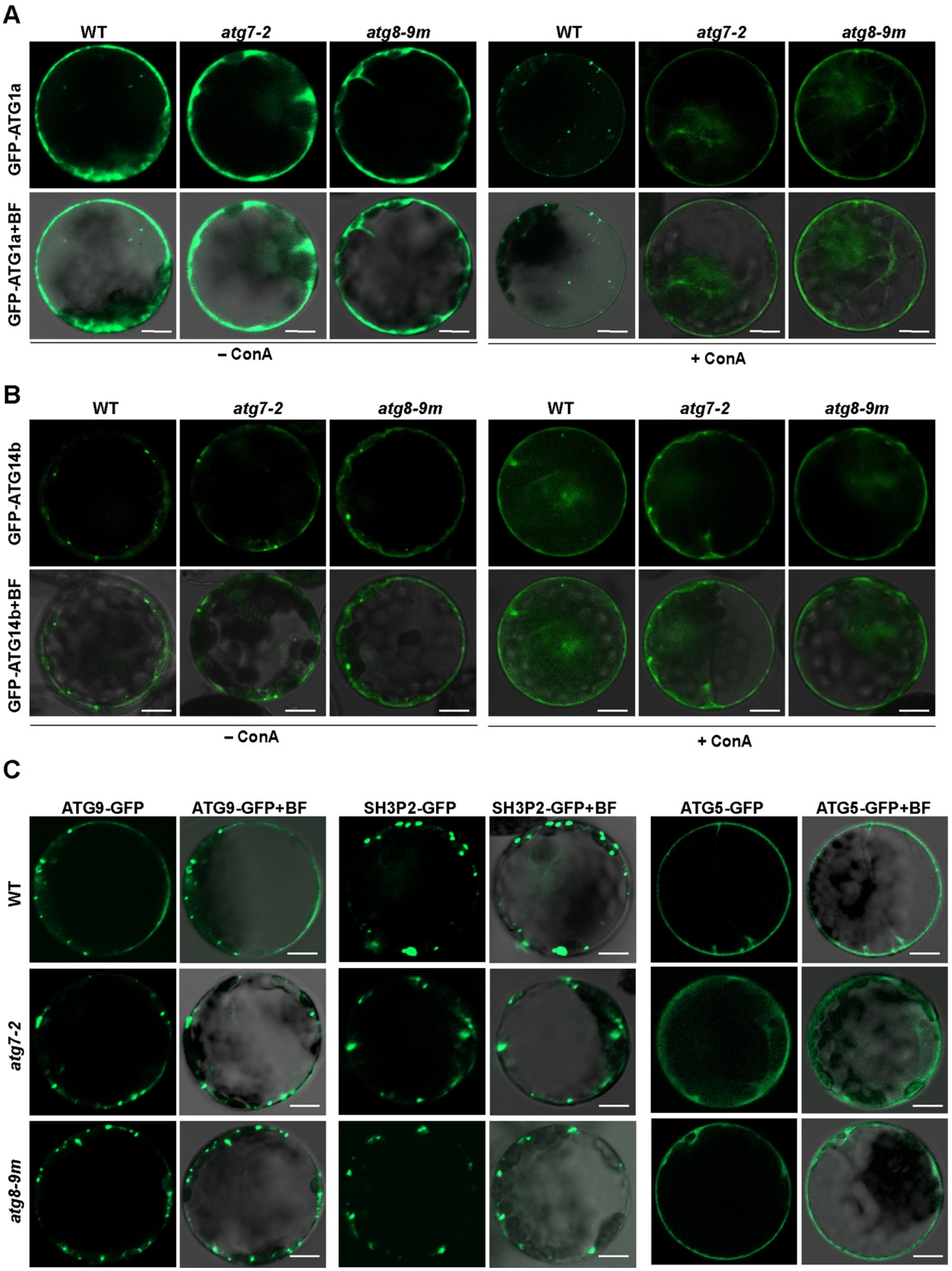


**Supplemental Figure 2.** Transient Expression of Different GFP-Fused ATG Proteins in *atg8-9m* Protoplasts.

Leaf protoplasts of WT, *atg7-2*, and *atg8-9m* mutants were transformed with different GFP-ATG constructs and analyzed by confocal microscopy 12 to 14 h after transfection. 1 μM ConA or DMSO (–ConA) was added to protoplasts transformed with GFP-ATG1a or GFP-ATG14b constructs. Bars = 10 μm.

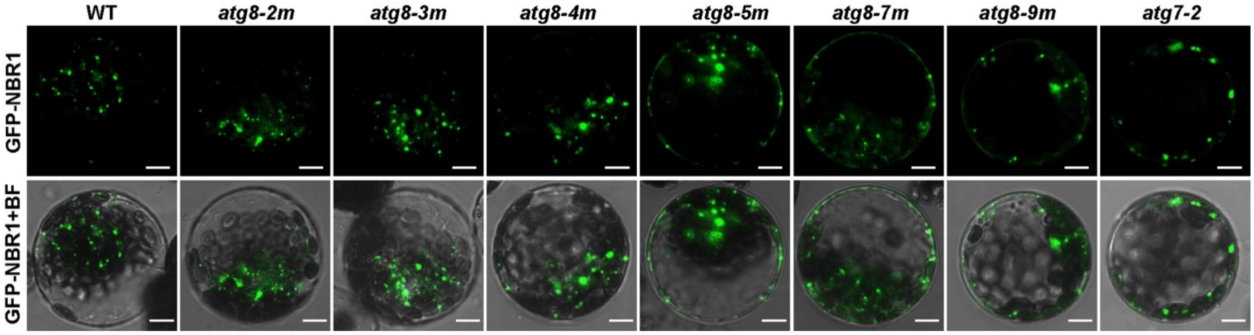


**Supplemental Figure 3.** Transient Expression of the GFP-NBR1 Fusion Protein in Protoplasts of Various *atg8* Mutants.

Leaf protoplasts of WT, *atg7-2*, and various *atg8* mutants were transformed with the GFP-NBR1 construct and then analyzed by confocal microscopy 12 to 14 h after transfection. Bars = 10 μm.

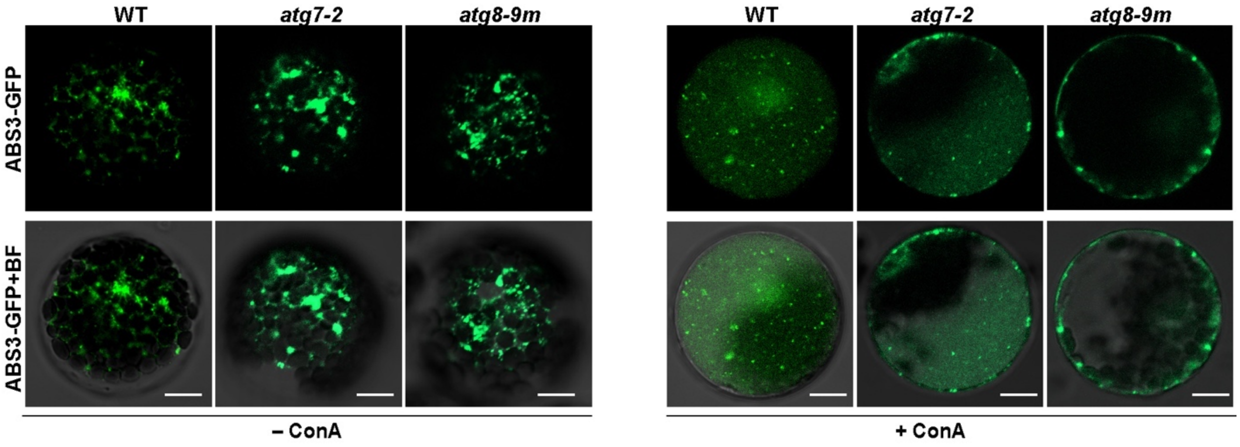


**Supplemental Figure 4.** Transient Expression of GFP-Fused ABS3 Protein in *atg8-9m* Protoplasts.

Leaf protoplasts of WT, *atg7-2*, and *atg8-9m* mutants were transformed with ABS3-GFP constructs and treated with 1 μM ConA or DMSO (–ConA) for 12 to 14 h prior to confocal imaging analysis. Bars = 10 μm.

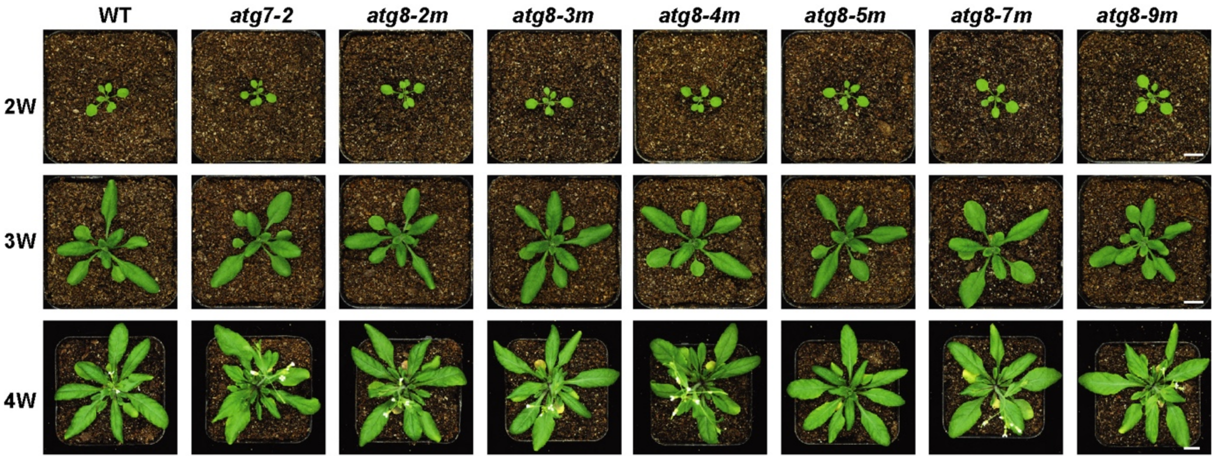


**Supplemental Figure 5.** Phenotypic Analysis of *Arabidopsis atg8* Mutants.

The wild-type Col-0 (WT), the autophagy mutant *atg7-2*, and various *atg8* mutants were grown on soil at 22°C under LD conditions. Photographs were taken at 2, 3, and 4 weeks after germination. Bar = 1 cm.

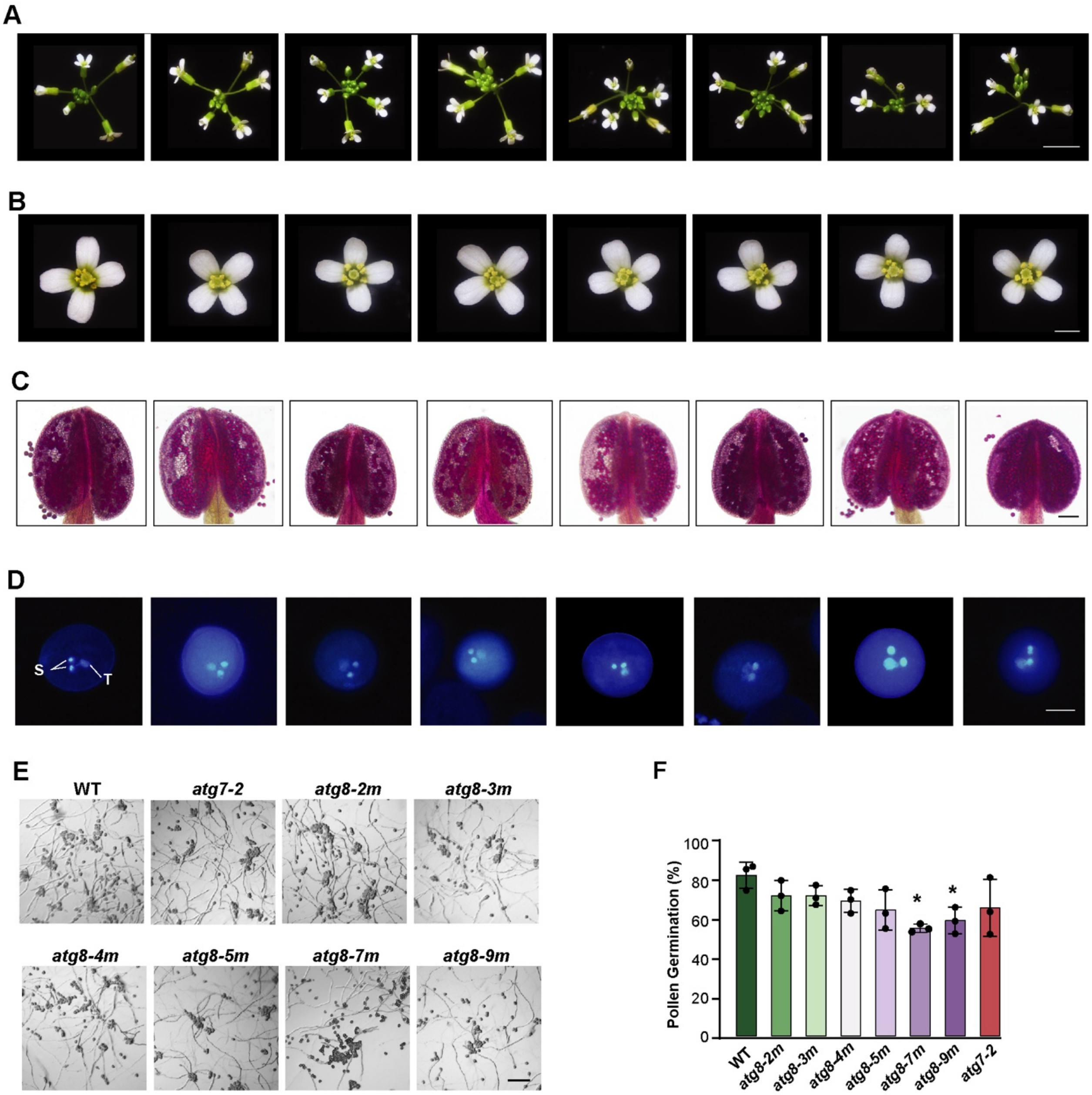


**Supplemental Figure 6.** *atg8* Mutants Produce Viable Pollen.

**(A-B)** Morphological observation of the inflorescence development of WT, *atg7-2*, and various *atg8* mutants grown under LD photoperiod for seven weeks. Bars = 5 cm (A) and 1 mm (B), respectively.

**(C)** Alexander staining of mature pollen grains. Bar = 1.0 mm.

**(D)** 4',6-diamidino-1-phenylindole (DAPI) staining to localize the tube nucleus (T) and two sperm nuclei (S) in mature pollen. Bar = 20 µm.

**(E-F)** *In vitro* pollen germination assays for *atg*8 mutants. (E) Representative photomicrographs of pollen germination on germination medium for 24 h. Bar = 0.5 mm. (I) Pollen tube germination rates. Different letters indicate significant differences ($p$ < 0.05) as determined by two-way ANOVA followed by Tukey's multiple comparison test. Three independent experiments were performed (*n* = 300 grains from ten anthers per genotype in each independent experiment).

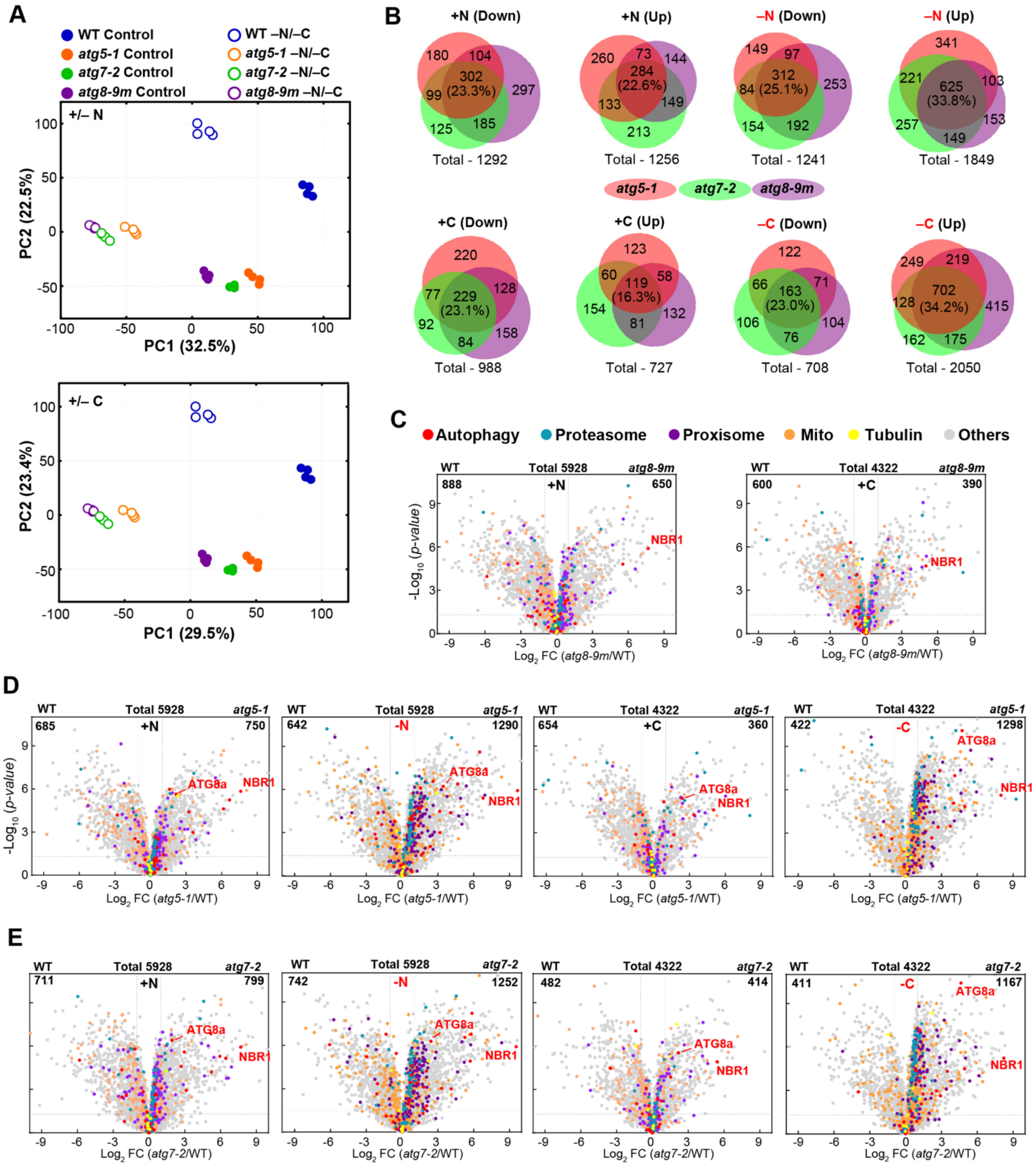

**Supplemental Figure 7**. Proteomic Comparisons of Wild Type and *atg* Mutants During N- or C-starvation.

**(A)** PCA of proteomic data sets for WT and *atg* mutants. Values were determined from $log_2$ transformed MS1 precursor ion intensities for the proteomic data sets (*n* = 4 biological replicates, each with two technical replicates). The amounts of variation explained by the first two components are indicated on the axes, and the numbers represent the PC scores.

**(B)** Venn diagram showing the overlap of proteins significantly affected by *atg* mutations compared to wild type under nitrogen or fixed-carbon stress.
**(C)** Volcano plots showing the preferential accumulation of proteins categorized by Go to specific processes, protein complexes, and cellular compartments in *atg8-9m* compared to wild-type seedlings under control growth conditions.
**(D-E)** Volcano plots showing the preferential accumulation of proteins categorized by Go to specific processes, protein complexes and cellular compartments in *atg5-1* (D) or *atg7-2* (E) versus wild-type seedlings under nutrient stress.

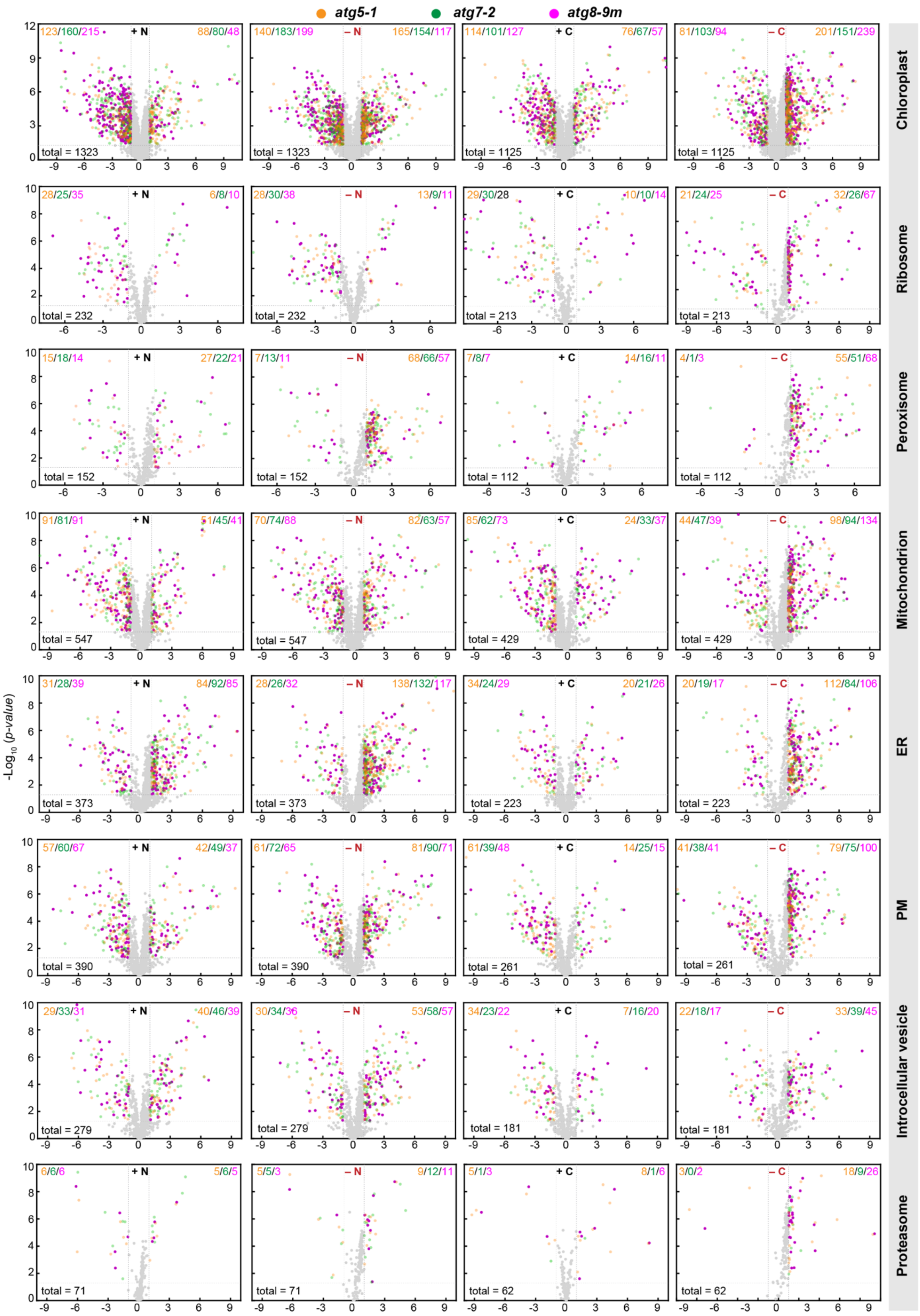

atg5-1
atg7-2
atg8-9m
+ N
– N
+ C
– C
-Log10 (p-value)
Chloroplast
Ribosome
Peroxisome
Mitochondrion
ER
PM
Introcellular vesicle
Proteasome
total = 1323
total = 1125
total = 232
total = 213
total = 152
total = 112
total = 547
total = 429
total = 373
total = 223
total = 390
total = 261
total = 279
total = 181
total = 71
total = 62

**Supplemental Figure 8.** The Influence of *atg8* Mutation on Protein Abundances in Specific Cellular Compartments/Complexes in N- or C-starved *Arabidopsis* Samples.

Volcano plots showing the preferential accumulation of proteins from GO subcategories related to specific cellular compartments and the 26S proteasome complex from *atg* mutants treated with nitrogen (+/–N) or fixed-carbon (+/–C) stress. Protein abundances were measured as shown in Figure 3. Each protein in the GO subcategory was plotted based on its log2 fold change (FC) in abundance and its -log10 P value in significance comparing WT with *atg5-1* (in orange), *atg7-2* (in green), or *atg8-9m* (in mCherry).

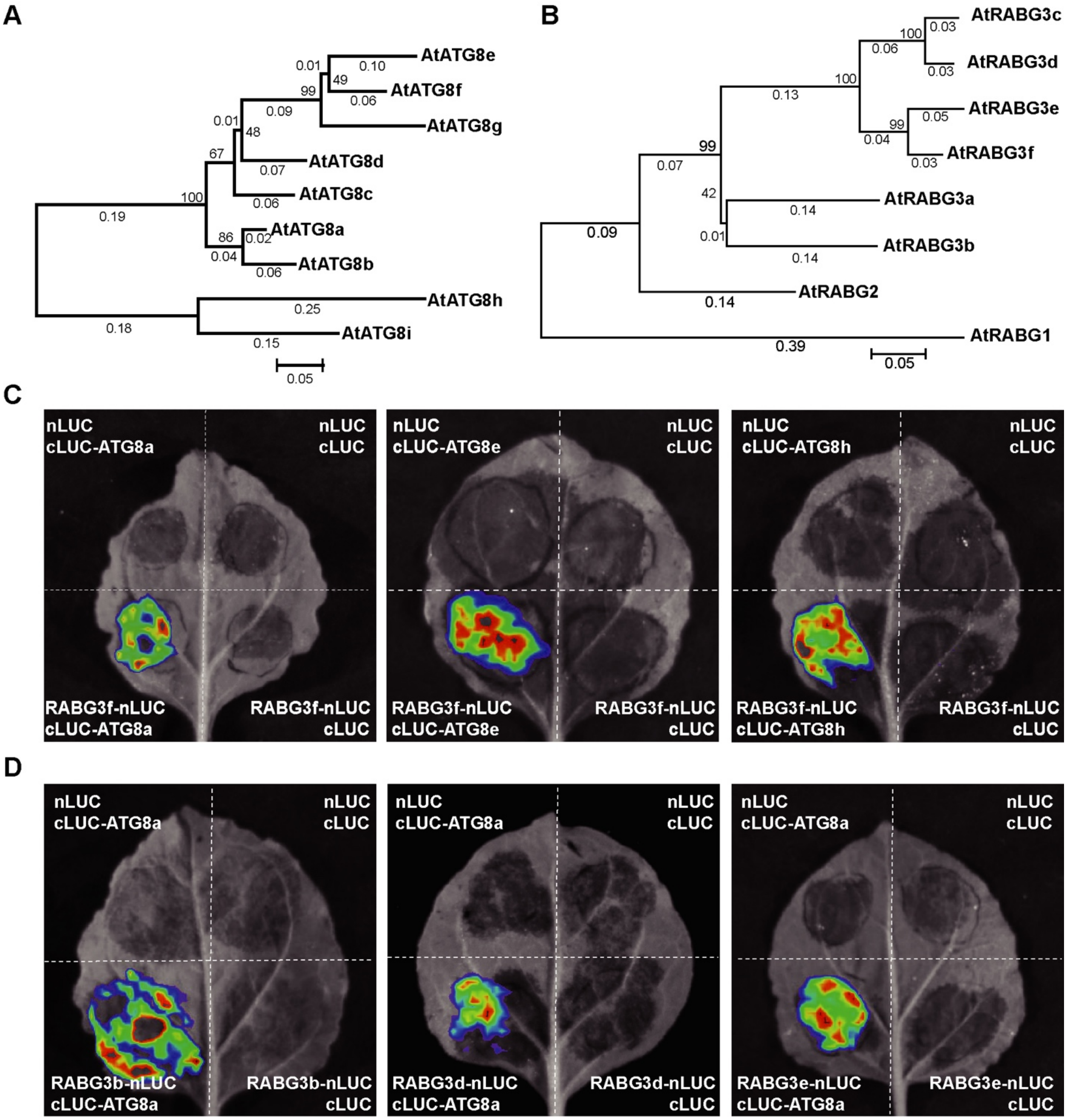


**Supplemental Figure 9.** LCI Assays for the Detection of RABG3-ATG8 Interactions.

**(A-B)** Phylogenetic analyses of *Arabidopsis* ATG8 (A) and RABG3 (B) proteins. The neighbor-joining phylogenetic trees were constructed with the MEGA v6.0 program using the full-length sequences of *Arabidopsis* ATG8 and RABG3 proteins. The percentage bootstrap values obtained from 1000 replicates are shown in the cladogram.

**(C)** LCI assays to detect the interactions between different ATG8 isoforms and RABG3f.

**(D)** LCI assays to detect the interactions between different RABG3 isoforms and ATG8a.

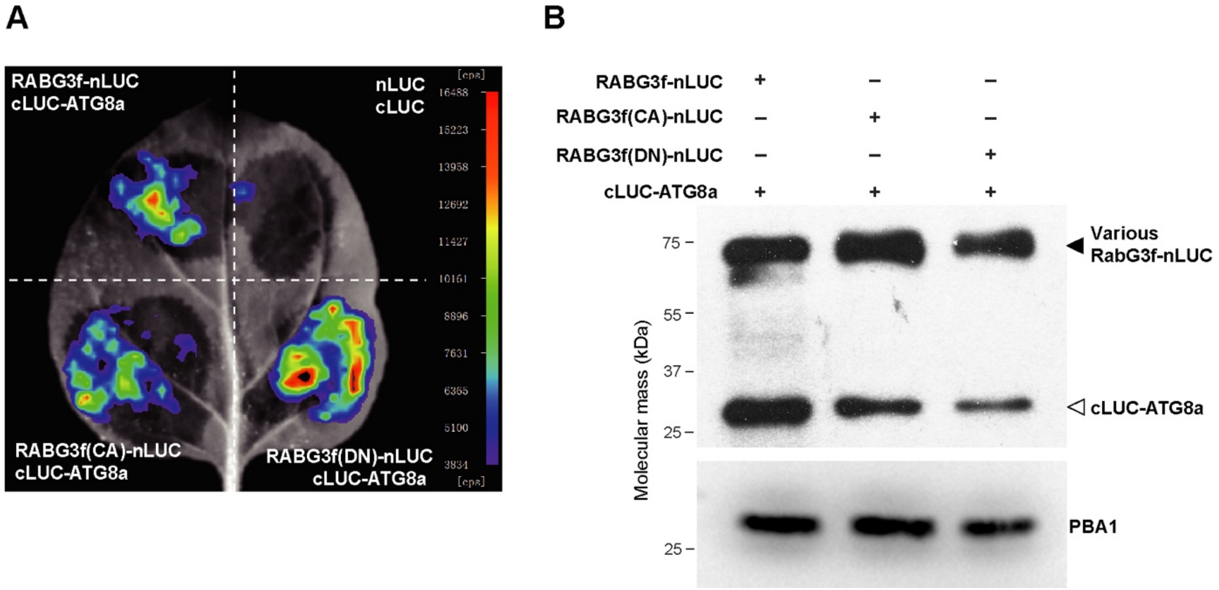


**Supplemental Figure 10.** Mutations in the GTPase Region of RABG3f Do Not Affect the RABG3f-ATG8a Interaction.

(A) LCI assays to detect the interactions between ATG8 and RABG3f in its wild-type, constitutively activated (CA), or dominant negative (DN) forms.

(B) Expression levels of cLUC and nLUC fusion proteins in (A) were validated by immunoblotting with anti-luciferase antibody which recognizes both nLUC and cLUC of luciferase. Antibody against PBA1 was used as a loading control.

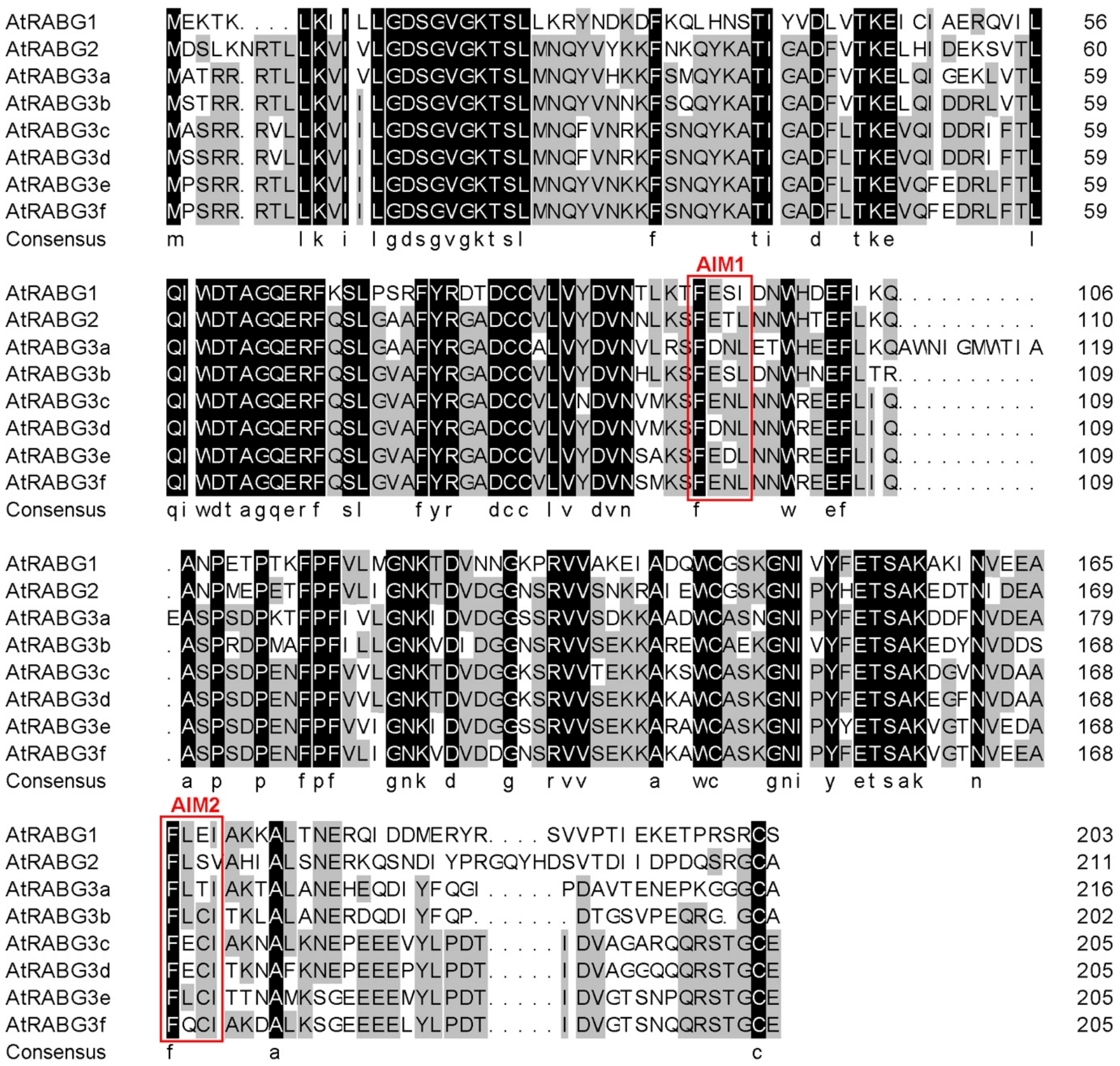


**Supplemental Figure 11.** Sequence Alignment of the *Arabidopsis* RABG Proteins.

Red boxes: potential AIMs predicted by hfAIM (http://bioinformatics.psb.ugent.be/hfAIM/).

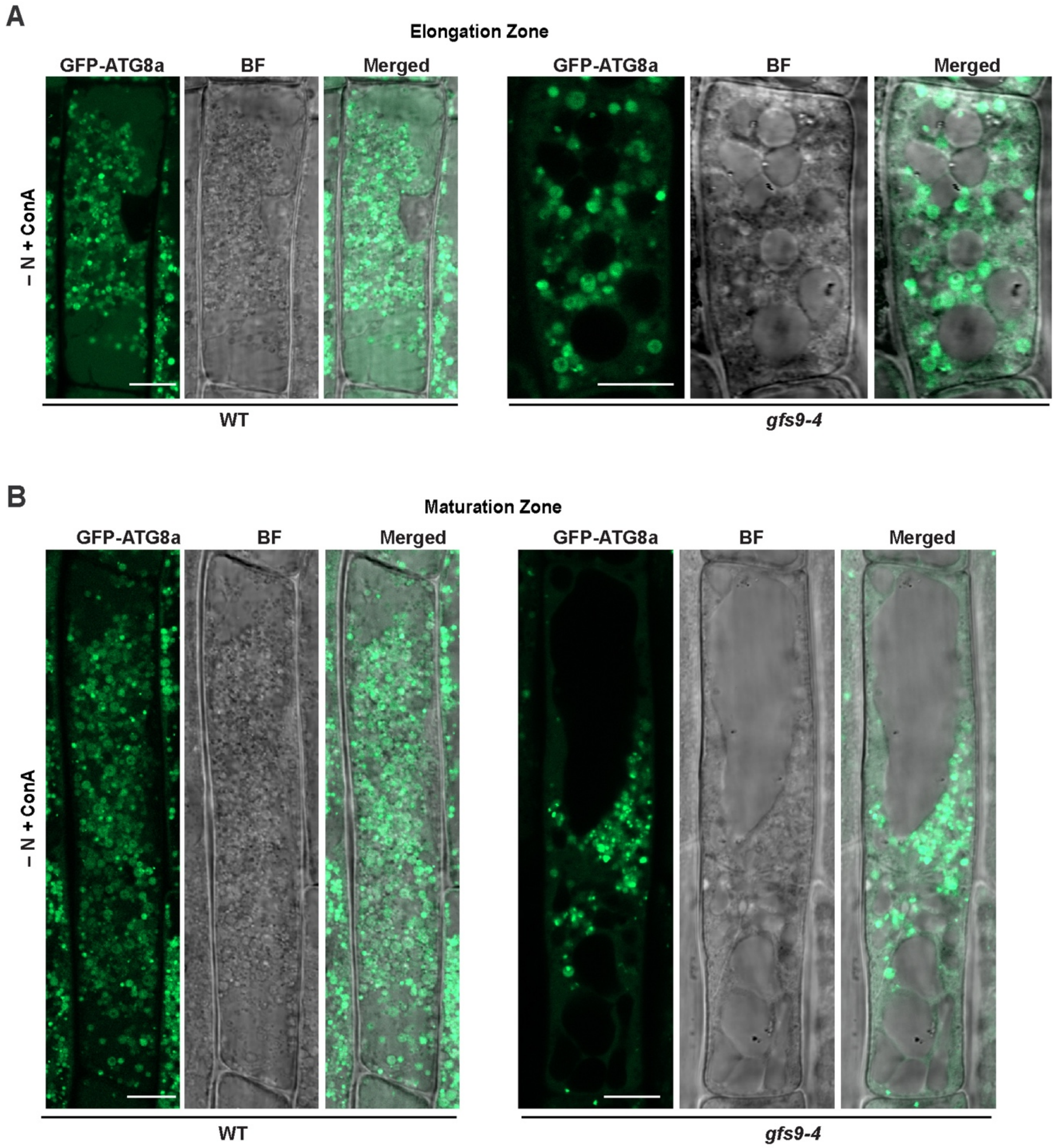


**Supplemental Figure 12.** Intracellular Distribution of the Autophagy Marker GFP-ATG8a in the Root Elongation and Maturation Zones of *gfs9-4* Seedlings.

Plants stably expressing the GFP-ATG8a marker were grown on MS medium for 6 d and then exposed to nitrogen-deficient medium containing 1 μM ConA for 12 h before confocal microscopy. Bars = 10 μm

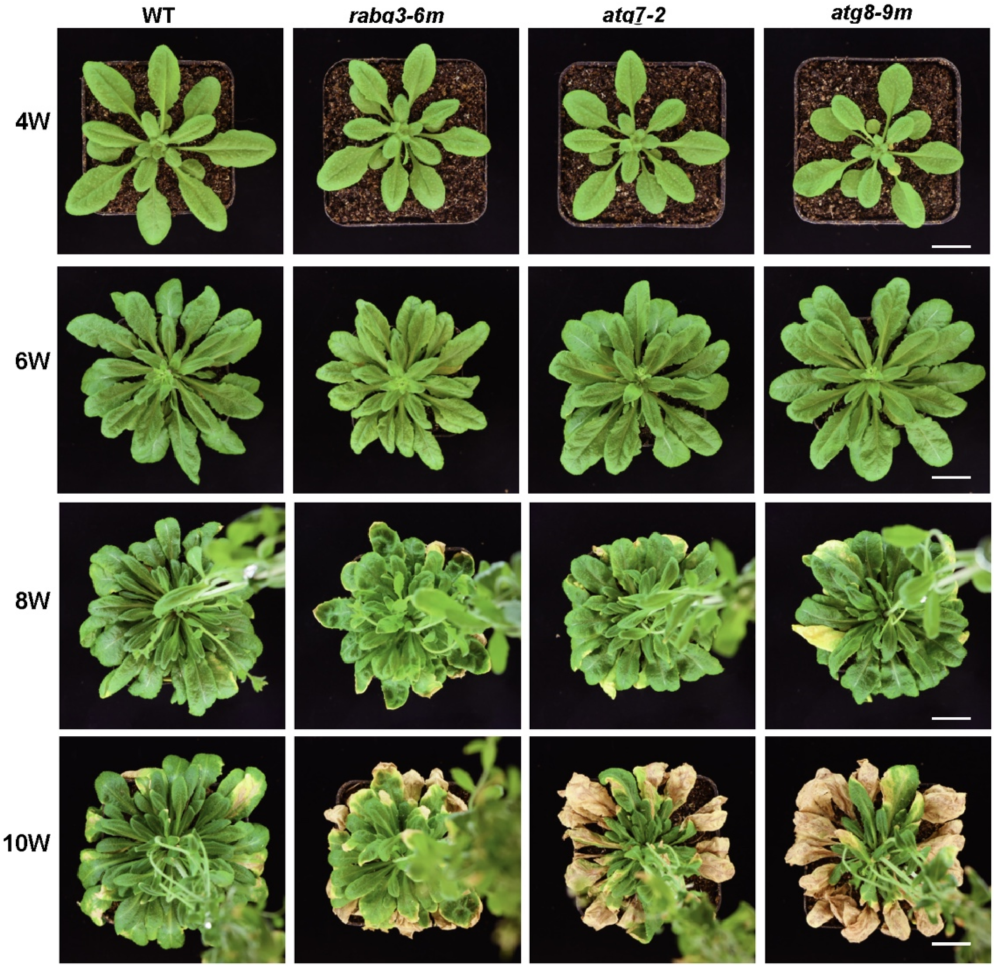


**Supplemental Figure 13.** The *rabg3-6m* Mutant Exhibited an Early Leaf Senescence Phenotype Under SD Conditions.

The wild-type Col-0 (WT), the homozygous *rabg3-6m* mutant, and the autophagy mutants *atg7-2* and *atg8-9m* were grown on soil at 22°C under SD conditions. Photographs were taken 4, 6, and 10 weeks after germination. Bar = 1 cm.

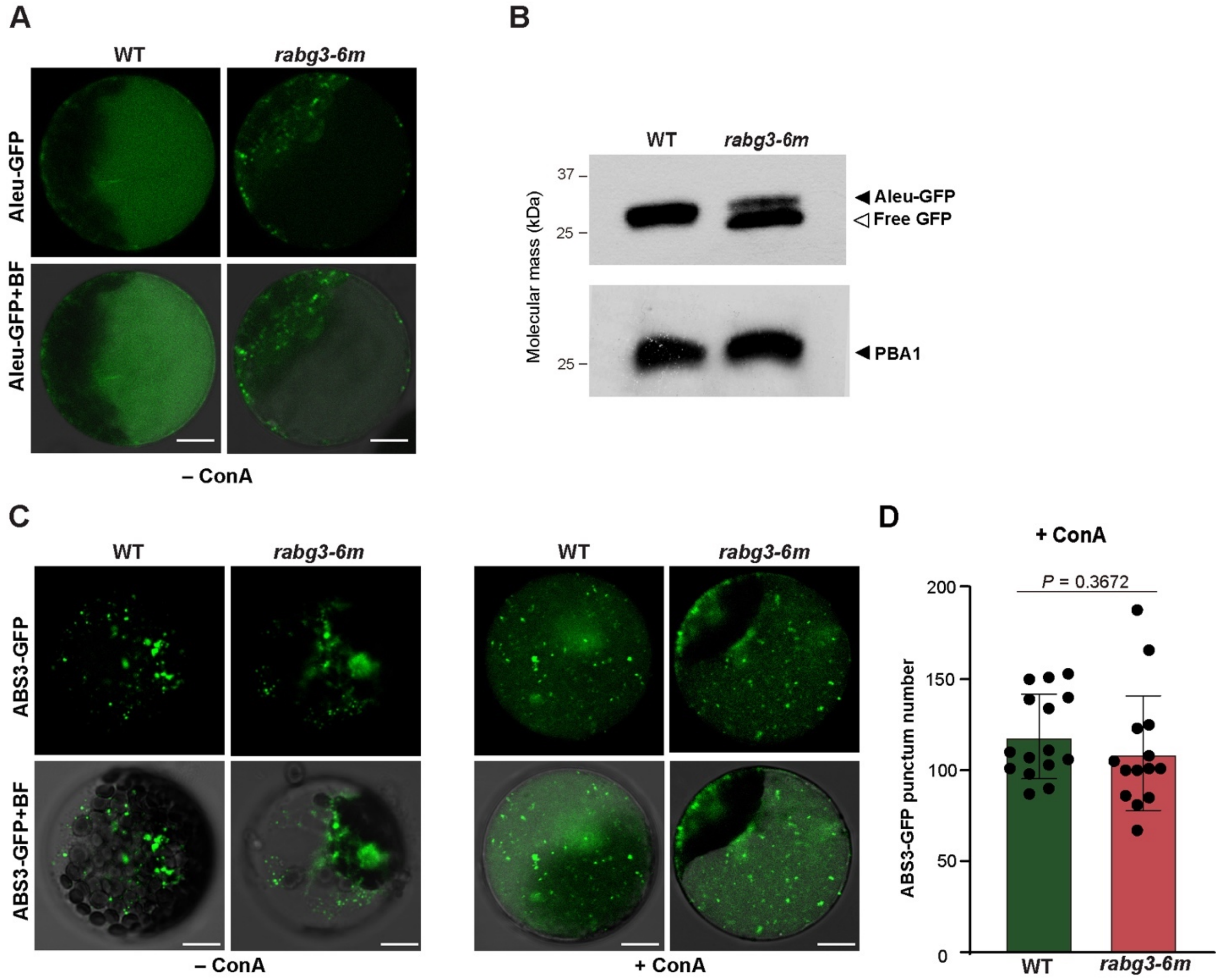


**Supplemental Figure 14.** RABG3 Is Required for the Vacuolar Deposition of Aleu-GFP But Not for ABS3-GFP.

**(A)** Vacuolar transport of Aleu-GFP in *rabg3-6m* protoplasts. Protoplasts of WT or *rabg3-6m* were transformed with the Aleu-GFP construct and incubated in liquid medium for 8 h prior to observation. Bars = 10 μm.

**(B)** Immunoblot analysis of *Arabidopsis* leaf protoplasts transiently expressing Aleu-GFP as shown in (A) with anti-GFP antibodies. The Aleu-GFP and free GFP are indicated by closed and open arrowheads, respectively. PBA1 was used to confirm nearly equal protein loading.

**(C-D)** Vacuolar transport of ABS3-GFP in *rabg3-6m*. Protoplasts of WT or *rabg3-6m* were transformed with the ABS3-GFP construct and incubated in liquid medium containing DMSO (–ConA) or ConA for 16 h prior to confocal microscopy observation. Bars = 10 μm. Quantification of vacuolar puncta of ABS3-GFP shown in (D). Images ($n$ = 15) were collected to measure the number of puncta. Statistical significance was determined by Student's *t*-test.

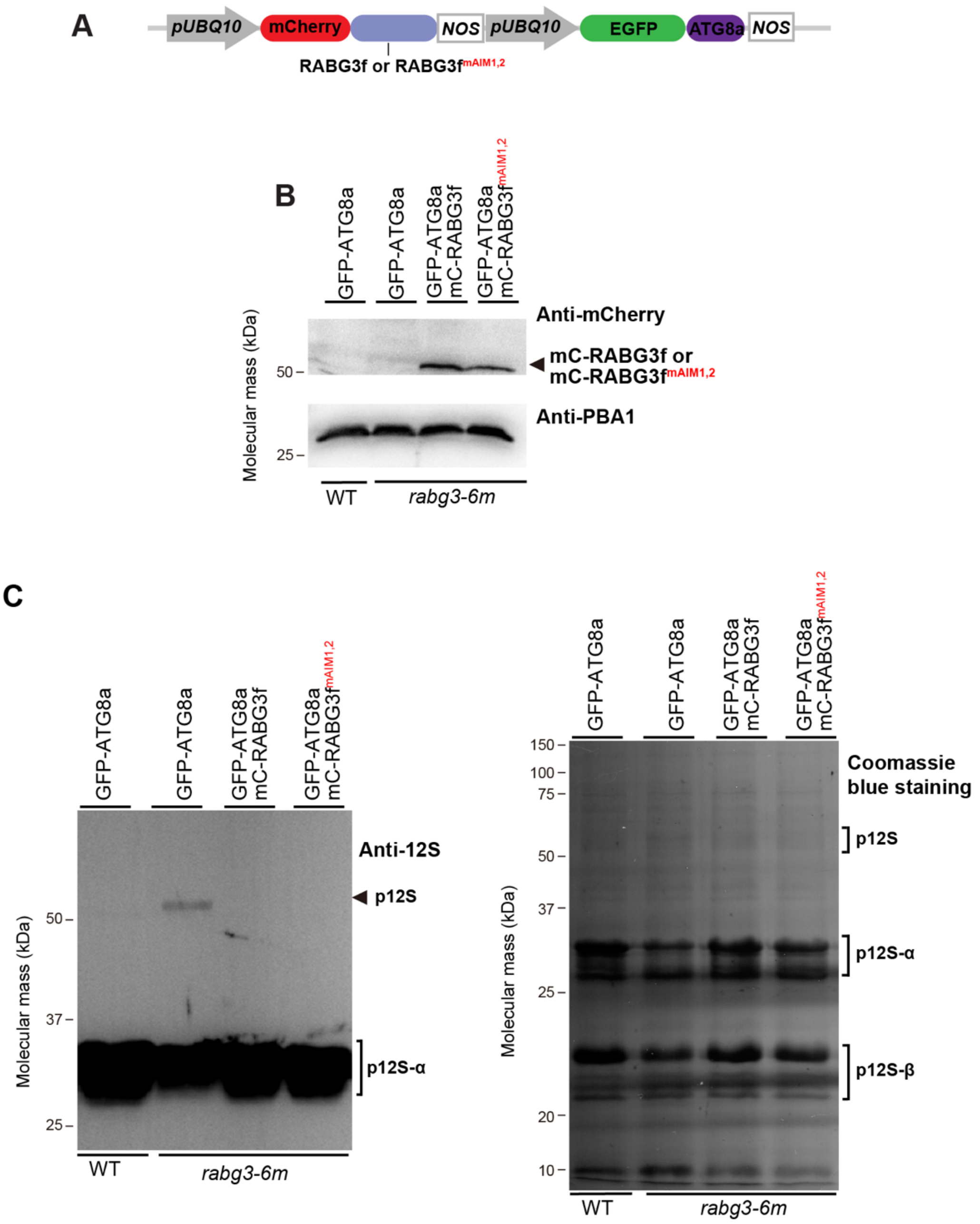


**Supplemental Figure 15.** Genetic Complementation of *Arabidopsis rabg3-6m* with Wild-Type or AIM-Mutated RABG3f.

**(A)** Schematic representation of the genetic constructs used for complementation analysis. The autophagy marker GFP-ATG8a is coexpressed with either wild-type or AIM-mutated RABG3f.

**(B)** Immunoblot analysis of complementation lines (GFP-ATG8a mCherry-RABG3f/*rabg3-6m* and GFP-ATG8a mCherry-RABG3f$^{mAIM1,2}$/*rabg3-6m*). Immunoblotting was performed with total proteins extracted from seven-day-old seedlings, and antibodies against mCherry were used to mCherry fused proteins. PBA1 was used as a loading control.

**(C)** Immunoblot analysis of complementation lines using anti-12S globulin antibodies. Immunoblotting was performed with total proteins extracted from dry seeds of the indicated genotypes. p12S, 12S globulin precursor; 12S-α and 12S-β, α and β subunits of mature 12S globulins. The SDS-PAGE profile of total seed proteins of the indicated genotypes after Coomassie blue staining is shown in the right panel.
